# The genetic population structure of Lake Tanganyika’s *Lates* species flock, an endemic radiation of pelagic top predators

**DOI:** 10.1101/2021.04.23.441176

**Authors:** Jessica A. Rick, Julian Junker, Ismael A. Kimirei, Emmanuel A. Sweke, Julieth B. Mosille, Christian Dinkel, Salome Mwaiko, Ole Seehausen, Catherine E. Wagner

## Abstract

Understanding genetic connectivity plays a crucial role in species conservation decisions, and genetic connectivity is an important component of modern fisheries management. In this study, we investigated the population genetics of four endemic *Lates* species of Lake Tanganyika *(Lates stappersii, L. microlepis, L. mariae* and *L. angustifrons)* using reduced-representation genomic sequencing methods. We find the four species to be strongly differentiated from one another (mean interspecific F_ST_ = 0.665), with no evidence for contemporary admixture. We also find evidence for strong genetic structure within *L. mariae,* with the majority of individuals from the most southern sampling site forming a genetic group that is distinct from the individuals at other sampling sites. We find evidence for much weaker structure within the other three species (*L. stappersii, L. microlepis,* and *L. angustifrons).* Our ability to detect this weak structure despite small and unbalanced sample sizes and imprecise geographic sampling locations suggests the possibility for further structure undetected in our study. We call for further research into the origins of the genetic differentiation in these four species—particularly that of *L. mariae—*which may be important for conservation and management of this culturally and economically important clade of fish.

## Introduction

Life history traits—including reproduction, growth, and dispersal—strongly influence gene flow and the genetic structure of populations (Ellegren & Galtier 2016; Manier & Arnold 2006; Stearns 1992). Differences in life history traits can also explain why sympatric and ecologically similar taxa may differ in their genetic population structure (Peterman *et al*. 2015; Young *et al*. 2015). Understanding how life history traits like dispersal, reproduction, and growth, shape gene flow is therefore crucial to improving our understanding of local adaptation, speciation, persistence of populations, migration, and the spatial scale of effective management (Sunday *et al*. 2014).

In teleost fishes, life history traits such as spawning location, philopatry, timing of spawning, and duration of larval stage, have been shown to shape population structure (Olsen *et al*. 2011; Pettersson *et al*. 2019; Young *et al*. 2015). Anadromous salmonids are well-known for displaying homing behavior that leads to genetic distinction between populations that use different spawning grounds at different times, dividing populations into many genetic units despite the populations spending most of their lives together in the ocean (Wenburg *et al*. 1998; Brannon *et al*. 2004). Divergence between populations caused by temporal differences in the timing of spawning has also been observed in Atlantic herrings (Martinez Barrio *et al*. 2016; Pettersson *et al*. 2019). In the oceans, coral reef fishes which spawn on or near the substrate, which exhibit parental care strategies, and which lack planktonic larvae, generally have significantly greater population structure than pelagic spawners or substrate spawners with planktonic larvae (Riginos *et al*. 2014). Additionally, a shorter larval phase among pelagic spawners can lead to a more structured population, because this leads to a shorter larval dispersal distance (Selkoe & Toonen 2011).

Understanding the genetic population structure of species is essential for conservation and sustainable fisheries management (Reiss *et al*. 2009; Bernatchez 2016; Supple & Shapiro 2018). Disregarding population structure and managing a population complex as a single population can lead to overfishing or even extinction of the more vulnerable genetic groups (Reiss *et al*. 2009; Sterner 2007). In species with cryptic diversity, unintentional overfishing of distinct populations can lead to a loss of genetic diversity in the species as a whole. Stock diversity can contribute to the resilience of a fishery in the face of environmental fluctuations (Schindler *et al*. 2015) and the loss of differentiated populations can result in major losses in the consistency of fisheries yields (Hutchinson 2008; Schindler *et al*. 2010). The loss of population diversity can also have dramatic effects on ecosystem services and species persistence; understanding the extent to which population structure exists is crucial for predicting these consequences (Schindler *et al*. 2015; Schindler *et al*. 2010; Therkildsen *et al*. 2013). It is therefore essential to understand the extent and nature of population genetic structure and differential adaptation in fishes that are important food and economic resources.

### The pelagic fish community and the fishery of Lake Tanganyika

Lake Tanganyika is the second largest inland fishery on the continent of Africa (Van der Knaap *et al*. 2014). During the last few decades, harvest records from Lake Tanganyika’s fishery have indicated a general decline in population sizes (Kimirei *et al*. 2008; Van der Knaap 2013; Van der Knaap *et al*. 2014; van Zwieten *et al*. 2002). This decline has resulted from a combination of an increase in the number of fishermen and vessels on Lake Tanganyika, changes in fishing practices (e.g., the use of beach seining, which targets fish in their in-shore nursery habitats) (Kimirei *et al*. 2008; Petit & Shipton 2012; Van der Knaap 2013; Van der Knaap *et al*. 2014; van Zwieten *et al*. 2002), and warming lake surface temperatures (Cohen *et al*. 2016; O’Reilly *et al*. 2003; Sayers *et al*. 2020). Decreases in fish abundance are likely also linked to reduced productivity in the lake caused by stronger water column stratification due to climate change (O’Reilly *et al*. 2003; Verburg *et al*. 2003). In addition, anthropogenic activity (e.g., land use changes, pollution, and increased sedimentation) impact both the water quality of Lake Tanganyika directly and the lake’s productivity via climate feedbacks (Ogutu-Ohwayo *et al*. 2016; Ivory *et al*. 2021; McGlue *et al*. 2021). Local fishermen similarly acknowledge spatial and temporal changes in fish abundance that are related to the ecological conditions of the lake (Bulengela *et al*. 2020; De Keyzer *et al*. 2020). Consequently, there is increasing recognition of the need to develop sustainable management strategies for the lake’s pelagic fish stocks (Kimirei *et al*. 2008; Mölsä *et al*. 1999; Mölsä *et al*. 2002; Van der Knaap 2013; Van der Knaap *et al*. 2014; van Zwieten *et al*. 2002).

Lake Tanganyika’s pelagic zone is relatively low in species diversity compared to that of other large African lakes: it consists predominantly of six endemic fish species belonging to two families (Coulter 1991). These include a monophyletic pair of mainly planktivorous clupeids *(Stolothrissa tanganicae, Limnothrissa miodon)* and four mainly piscivorous latids of the genus *Lates (L. stappersii, L. mariae, L. microlepis, L. angustifrons).* While all four *Lates* species likely have planktivorous larvae (van Zwieten *et al*. 2016), only *L. stappersii* remains pelagic throughout its life (Ellis 1978). The other three species move inshore to littoral weed beds as juveniles (>3cm) before *L. microlepis* returns to the pelagic, *L. mariae* transitions to the deepwater benthic zone, and *L. angustifrons* can be found throughout the water column as adults (Coulter 1991). Of the four *Lates* species, *L. stappersii* is the most abundant and is an important component of the current fishery of Lake Tanganyika (Coulter 1976, 1991; Kimirei *et al*. 2008; Mannini 1998a, b; Mölsä *et al*. 1999; Mölsä *et al*. 2002; Munyandero 2002; Sarvala *et al*. 2002; Van der Knaap 2013; Van der Knaap *et al*. 2014; van Zwieten *et al*. 2002).

The fish of Lake Tanganyika are influenced by a complex interplay of bottom-up control via different mixing regimes and upwelling rates (Bergamino *et al*. 2010; O’Reilly *et al*. 2003; Stenuite *et al*. 2007; Verburg & Hecky 2009; Verburg *et al*. 2003), and top-down control via predators and fishing (Kimirei *et al*. 2008; Mannini 1998a; Mannini *et al*. 1996; Munyandero 2002; Van der Knaap 2013; Van der Knaap *et al*. 2014; van Zwieten *et al*. 2002). If spatially heterogeneous environmental factors, such as mixing regimes and nutrient supply, are consistent over generations and coupled with limited gene flow between spatially segregated populations, divergent selection on life history traits (e.g., spawning phenology, developmental timing and recruitment success) of fish in different parts of the lake could develop and persist. Spatial environmental variation in Lake Tanganyika, in combination with its large geographic extent, might therefore generate intraspecific genetic differentiation among populations of pelagic fish, as has been observed in pelagic cichlids in both Lake Tanganyika and Lake Malawi (Genner *et al*. 2010; Koblmüller *et al*. 2019). In these cichlids, strong spawning site fidelity (Genner *et al*. 2010) and differences in foraging behavior (Koblmüller *et al*. 2019) influence spatial genetic structure. In contrast, recent studies have found that the two pelagic clupeids in Lake Tanganyika show no evidence of spatial genetic structure (Junker *et al*. 2020; De Keyzer *et al*. 2019; Kmentová *et al*. 2020). Intraspecific spatial heterogeneity in spawning times (Coulter 1976; Ellis 1978), dominant diet items (Mannini *et al*. 1999), sizes at maturity (Mannini et al. 1996), and catch rates (Coulter 1991), along the length of Lake Tanganyika support the possibility that gene flow may be reduced among fish in different parts of the lake. Indeed, in *L. stappersii,* a study based on RAPD markers found individuals from the Kigoma region to be genetically differentiated from the rest of the sites sampled in this species (Kuusipalo 1999), with some additional differentiation observed among individuals collected during the dry versus rainy.

Here, we use reduced-representation genomic data to test the prediction that differences in life history traits among the four Lake Tanganyika-endemic *Lates* species result in differences among the species in their intraspecific genetic connectivity and population structure. We first assessed whether phenotypic identification coincides with genetically based species boundaries and then examined the extent to which each of the four species is genetically structured, as well as how the genetic structure observed in each species corresponds to its known life history and the environment. Based on known differences in life history among the four species, we hypothesized that we would find little genetic population structure in the entirely pelagic *L. stappersii* and mainly pelagic *L. microlepis.* In contrast, we expected the highest genetic differentiation in populations of the predominantly benthic *L. mariae.* Finally, we hypothesized that *L. angustifrons* would show intermediate amounts of structure since there is no evidence for large scale movements of this species, even though they are found throughout the water column.

## Material and Methods

### Study system and sampling

Between 2001 and 2019, we collected tissue samples (fin clips) from *L. stappersii, L. mariae, L. microlepis,* and *L. angustifrons* at 9 general locations along the Tanzanian shore of Lake Tanganyika, which together span the ~490 km shoreline from the northern to southern border (Fig. 1; Table S2). All fish collections were made in partnership with researchers of the Tanzanian Fisheries Research Institute (TAFIRI) in Kigoma, Tanzania; most of the fish were obtained opportunistically from fishermen, and therefore were already dead when we received them. Given fuel costs and limitations of most outboard motors used by fishermen in this region, fishermen generally fish within a 20km radius of their landing sites (personal observation), but there remains inherent uncertainty in the exact location and depth at which these fish were collected. Fish were identified in the field by sampling personnel following United Nations Food and Agricultural Organization (FAO) identification guidelines (Eccles 1992) and guidance from Tanzanian researchers and local fishermen, and genetic methods were later used to confirm species identities for each fish. The majority of fish were collected during two targeted sampling campaigns in 2017 and 2018, the first of which occurred at the end of the dry season (September-October 2017), while the second occurred just following the rainy season (April-May 2018). For fish collected from 2016-2019, we recorded length and weight, and took standardized pictures of each fish. For the few live fish that were caught as part of other research, we took cuvette photographs of the live fish and subsequently euthanized the fish with an overdose of MS222. We then took fin clips for genetic analysis from all fish. Specimens smaller than 600mm were preserved in formaldehyde and archived in the collections at TAFIRI (Kigoma, Tanzania), EAWAG (Kastanienbaum, Switzerland), the University of Wyoming Museum of Vertebrates (Laramie, WY, USA), or the Cornell University Museum of Vertebrates (Ithaca, NY, USA).

**Figure 1.**
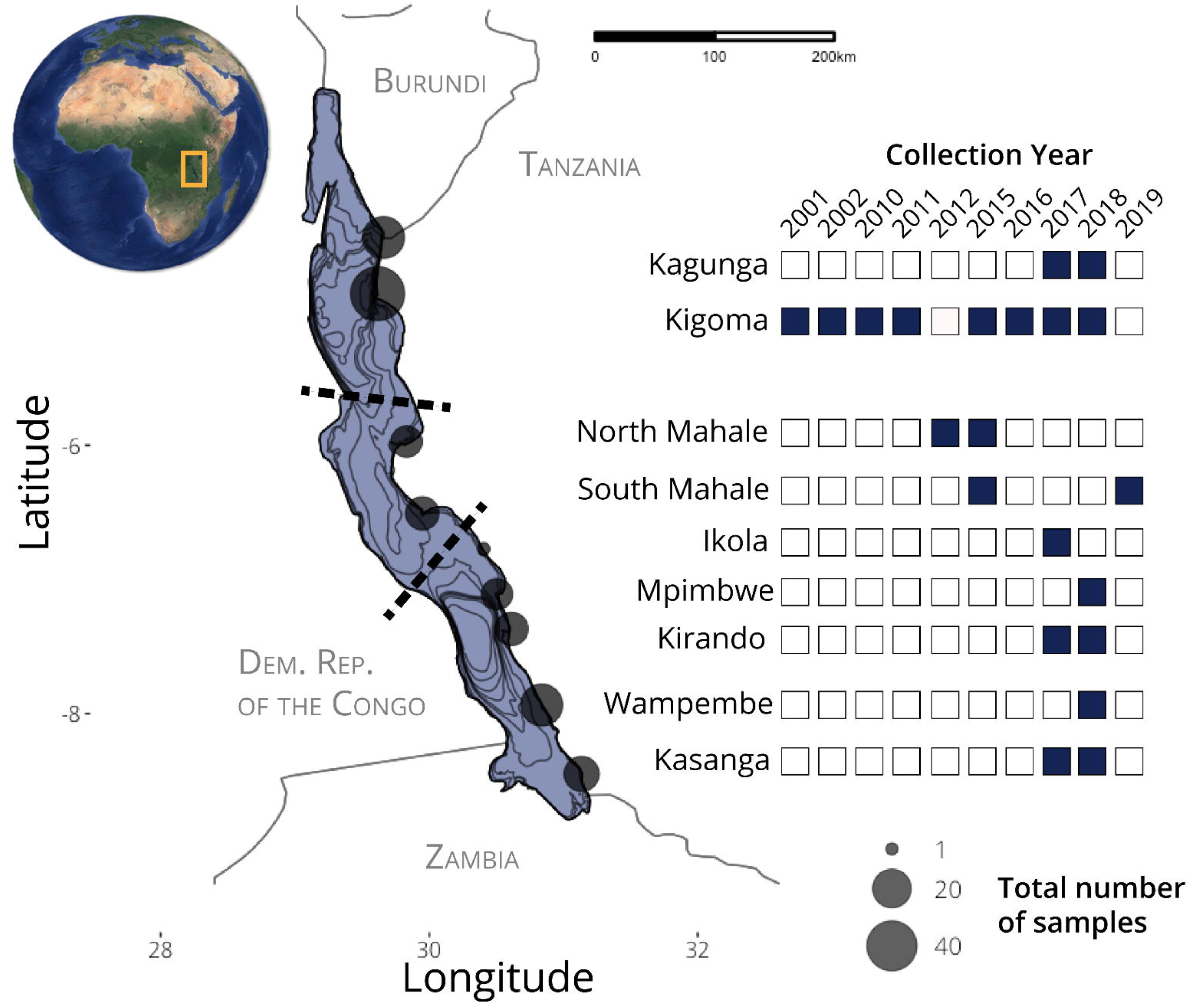
Sampling site locations and years in which fish were collected at each location (shaded boxes). Circle sizes on the map are proportional to the number of samples in the final genotyping-by-sequencing dataset for all species combined, with a minimum of 1 and maximum of 45 samples collected per site across all years. Dashed lines indicate the general boundaries of the north, central, and southern basin boundaries within the lake. Additional sampling information is available in Table S1 and Table S2.

### Genomic sequencing

We extracted DNA from fin clips using DNeasy Blood and Tissue Kits (Qiagen, Inc.) following the standard protocol, with the addition of an RNAse A incubation step. We then prepared genomic libraries for a combination of genotyping-by-sequencing (GBS, for *L. stappersii, L. mariae, L. microlepis, L. angustifrons)* and restriction site associated DNA sequencing (RAD, for *L. stappersii).*

We prepared GBS libraries following protocols outlined in Parchman *et al*. (2012), using MseI and EcoRI restriction enzymes. Following fragmentation via restriction enzyme digestion, fragmented DNA was barcoded by ligating short, unique individual-identifying DNA fragments (barcodes) to each individual’s fragmented DNA. These barcoded fragments were then amplified by PCR and pooled for sequencing. We ran two replicate PCRs for each individual and pooled the final PCR products into two libraries. The prepared libraries were size-selected for 200-350bp fragments using Blue Pippin (Sage Science, MA). One library was sequenced on one lane of Illumina HiSeq4000 (150bp single-end) at the University of Texas at Austin’s Genome Sequencing and Analysis Facility (UT GSAF; Austin, TX) and the other library was sequenced on one lane of Illumina HiSeq4000 (150bp, single-end) at the University of Oregon’s Genomics and Cell Characterization Core Facility (Eugene, OR). Each library contained 192 individuals, and select individuals were duplicated across libraries to check for library compatibility.

The RAD libraries containing *L. stappersii* were prepared for sequencing following the protocol by Baird *et al*. (2008) with the following modifications: we used between 400ng and 1000ng genomic DNA per sample and digested with *SbfI* overnight. We tagged each individual using P1 adapters (synthesized by Microsynth) with custom six to eight base pair barcodes and multiplexed 67 barcoded individuals per library, resulting in a total of three RAD libraries. The libraries were sheared using an S220 series Adaptive Focused Acoustic (AFA) ultra-sonicator (Covaris) with the manufacturer’s settings for a 400 bp mean fragment size. We size-selected for fragments between 300 and 700bp using a sageElf (Sage Scientific Electrophoretic Lateral Fractionator; Sage Science, Beverly, MA). The enrichment step was done in 6 aliquots with a total volume of 200 μl. Volumes were combined prior to the final size selection (again 300-700bp) step using the sageELF. Sequencing was done by the Lausanne Genomic Technologies sequencing facilities (University of Lausanne, Switzerland). All three RAD libraries were single-end sequenced on one lane each of the Illumina HiSeq2000 (100bp SE).

### Sequence data preparation

We filtered raw sequencing reads from each library by first removing the reads derived from the PhiX genome and other common contaminants (e.g. excess barcodes, primers, and adapters) using bowtie2 (v2.0.0, Langmead & Salzberg 2012). We filtered reads for an intact enzyme restriction site *(SbfI* for RAD libraries, MseI and EcoRI for GBS libraries), demultiplexed the fastq files, and matched sequence barcodes to individual fish using custom perl and bash scripts.

For the GBS libraries, the reads were mapped to the chromosome-level *Lates calcarifer* reference genome (v3, NCBI Genome Assembly GCA_001640805.1; Vij *et al*. 2016) using bwa mem (v0.7.17; Li & Durbin 2009) with default settings. Following alignment, we excluded any individual with < 50,000 reads or less than 60% of raw reads assembled to the reference genome. We identified variable sites (i.e. single nucleotide polymorphisms; SNPs) in the assembly using SAMtools mpileup (v1.8; Li *et al*. 2009) and bcftools (v1.8; Li *et al*. 2009). In calling variable sites, we omitted indels and kept only high-quality biallelic variant sites (QUAL > 20 and GQ > 9). We then filtered SNPs by minor allele frequency and amount of missing data using vcftools (Danecek *et al*. 2011), additionally only calling genotypes with a minimum read depth of 5. We then selected only SNPs with a minor allele frequency greater than 0.01 that were present in at least 50% of individuals. Finally, we thinned each data set to retain one SNP per locus (-thin 90). We assessed the distribution of missing data across individuals and mean read depth across sites using vcftools (Danecek *et al*. 2011), additionally using whoa (v0.0.2.999, Anderson 2018) in R (v4.0.4, R Core Team 2021) to assess heterozygote miscall rate at low read depth sites to ensure that our read depth filter was stringent enough.

For the RAD libraries, we barcode-trimmed reads down to 84 nucleotides using process_radtags from Stacks (Catchen *et al*. 2013). The FASTX-toolkit v.0.0.13 (http://hannonlab.cshl.edu/fastx_toolkit/) was used for quality filtering. In a first step, we kept only reads with all base quality scores greater than 10; in a second step, we removed all reads with more than 5% of the bases with quality score below 30. The RAD library was then mapped to the *Lates calcarifer* reference genome (Vij et al. 2016) and filtered, following the same steps as for the GBS libraries above (MAF > 0.01, missing data < 0.5, one SNP per locus, read depth > 5). There was no overlap of individuals included in the RAD and GBS sequencing libraries.

After filtering variants, we calculated the distribution of minor allele reads at heterozygous sites for each individual in both the RAD and GBS libraries, using a custom bash script (available at http://github.com/jessicarick/lates-popgen). If there is an excess of sequencing errors or contamination for an individual, then we would expect the ratio of minor allele to major allele reads to be significantly less than 1:1. Thus, we plotted the distribution of minor allele reads at heterozygous sites for each individual and removed individuals with a deficiency in minor allele reads across the majority of heterozygous sites. After removing these individuals, we then re-called genotypes and re-filtered the SNP data set to produce our final SNP data set.

### Species assignment

Not all samples could be identified to species in the field. Some fish were caught in deep water and brought quickly to the surface by the fishermen, making them difficult to identify due to barotrauma. Furthermore, juveniles of the large species *L. microlepis, L. mariae* and *L. angustifrons* can be difficult to distinguish. We used three different lines of evidence to confirm species identification and ensure that morphological identification matched genetic assignment: (1) phenotypic identification based on FAO species descriptions; (2) individual ancestry assignment using a model-based clustering approach; and (3) inference of monophyletic groups in phylogenetic analysis. The details for the latter two of these steps are described below.

With the GBS data, we first visualized clusters in our data using principal component analysis using EMU (Meisner *et al*. 2021). EMU uses an iterative method to infer population structure in the presence of missing data and is therefore able to infer population structure even for datasets with high, uneven, or non-random proportions of missing data, which tend to produce bias in other PCA methods (Meisner *et al*. 2021). We used the EM acceleration method in EMU, using 10 eigenvectors for optimization (-e 10) with a maximum of 100 optimization iterations and keeping all eigenvectors as output. We then inferred species groups using the model-based genetic clustering program entropy (Gompert *et al*. 2014; Shastry *et al*. 2021). Entropy infers population structure from multilocus SNP data by assigning individuals partially or completely to groups using Bayesian estimation from genotype likelihoods. In taking uncertainty about individual genotypes into account via genotype likelihoods, the model integrates outcomes over genotype uncertainty. We converted our VCF containing sites with <50% missing data to the mpgl genotype likelihood format using the vcf2mpgl R script (available at http://github.com/iessicarick/lates-popgen; adapted from https://bitbucket.org/buerklelab/mixedploidy-entropy/src, v2.0). We then ran entropy for K=4 clusters, running three independent MCMC chains of 80,000 total steps, discarding the first 10,000 steps as burn-in, and retaining every 10^th^ value (thin=10), resulting in 7000 samples from the posterior distribution of each chain. Our MCMC chains were run in parallel using GNU parallel (Tange 2011). We checked MCMC chains for mixing and convergence of parameter estimates by plotting a trace of the MCMC steps. We then visualized assignments in R and assigned individuals to species based on group assignment probabilities (q).

We further corroborated species identities using maximum likelihood phylogenetic inference methods in RAxML (v8.1.17; Stamatakis 2014). We concatenated all SNPs, removed invariant sites using the raxml_ascbias python script (v1.0, from https://github.com/btmartin721/raxml_ascbias), and used the Lewis correction for invariant sites with the ASC_GTRGAMMA model of molecular evolution (Lewis 2001) within RAxML to infer the maximum likelihood phylogeny, including *L. calcarifer* (Vij et al. 2016) as the outgroup. We used 100 rapid bootstraps to estimate confidence in our maximum likelihood tree. We then visually identified monophyletic groups on our maximum likelihood tree and used these groupings to further verify species identities from entropy and PCA clusters.

### Genetic diversity

We calculated genetic diversity for each species in ANGSD (v0.931, Korneliussen *et al*. 2014). From the BAM alignment files, we first calculated the site allele frequency likelihoods based on individual genotype likelihoods (option -doSaf 1) using the samtools model (option – GL 1), with major and minor alleles inferred from genotype likelihoods (option -doMajorMinor 1) and allele frequencies estimated according to the major allele (option -doMaf 2). We filtered sites for a minimum read depth of 10, minimum mapping quality of 20, and minimum quality (*q*-score) of 20. From the site allele frequency spectra, we then calculated the maximum-likelihood estimate of the folded site frequency spectra (SFS) using the ANGSD realSFS program (with option -fold 1). The folded SFSs were then used to calculate genomewide genetic diversity (Watterson’s *θ*), using the ANGSD thetaStat program (Korneliussen *et al*. 2013).

### Population genetic structure and isolation-by-distance

For each species, we then re-called variants to create a species-specific set of SNPs from the GBS data. As with the species combined GBS data, we identified variable sites using SAMtools mpileup and bcftools, omitting indels and keeping only high-quality biallelic variant sites (QUAL > 20 and GQ > 9). We then filtered SNPs by minor allele frequency (> 0.01) and amount of missing data (<50% missing) using vcftools (Danecek et al. 2011), additionally only calling genotypes with a minimum read depth of 5. Finally, we thinned to one SNP per locus (-thin 90).

For each species, we then performed principal component analysis (PCA) using EMU (Meisner *et al*. 2021), after using PLINK (v1.90b6.9, Purcell *et al*. 2007; Chang *et al*. 2015) to convert our VCF into BED format. EMU is able to infer population structure even for datasets with high, uneven, or non-random rates of missing data, which tend to produce bias in other PCA methods (Meisner *et al*. 2021). We again used the EM acceleration method, using 10 eigenvectors for optimization (-e 10) with a maximum of 100 optimization iterations and keeping all eigenvectors as output. We also performed PCA in EMU on the RAD dataset for *L. stappersii.* Individuals that had been duplicated among libraries were kept separate through these initial analyses, and then their reads were combined and variants were recalled for each species after ensuring that the duplicates clustered near to one another in both species identification and intraspecific PCA analyses.

We estimated heterozygosity at each site using the dartR package in R (v1.9.9, Gruber *et al*. 2018). We then used the Reich-Patterson F_ST_ estimator (Reich *et al*. 2009), which is robust even with small sample sizes, to calculate genetic divergence between sampling sites within each species. We calculated Reich-Patterson F_ST_ estimates using a custom function in R (available at http://github.com/jessicarick/reich-fst), using functions from the vcfR (v1.12, Knaus & Grunwald 2017) and dartR packages for importing and manipulating our VCF. We then tested for isolation-by-distance (IBD) using a Mantel test (mantel.randtest from ade4 in R; v1.7-16, Dray & Dufour 2007) to calculate the correlation between the natural logarithm of Euclidean geographic distance and standardized genetic distance (F_ST_/1-F_ST_) for all pairwise combinations of sampling sites within each species. We chose to pool these tests by sampling site, as we do not have more specific locations for where most of the fish were caught.

To test for intraspecific population subdivision more formally, we again used entropy. For each species, we ran entropy for K=1 to K=6 to infer the most likely number of ancestral groups within each species, and to assign individuals to these clusters. We ran three independent MCMC chains of 100,000 total steps, discarding the first 10,000 steps as burnin, and retaining every 10^th^ value (thin=10), resulting in 9000 samples from the posterior distribution of each chain. We checked MCMC chains for mixing and convergence of parameter estimates by plotting a trace of the MCMC steps. We then calculated the deviance information criterion (DIC) for each value of K and used the model with the lowest DIC as our best for modeling the variation observed in our data. From our entropy results, we assigned individuals to intraspecific groups using a cutoff of q > 0.6 at each value of K. We then calculated the mean of Reich-Patterson F_ST_ estimates between these K groups to understand how genetically distinct these groups are from one another. We also ran entropy at K=2 and K=3 using datasets with a more stringent missing data filter (miss=0.9) to test whether group assignments were contingent on the choice of bioinformatic filters. We also calculated Reich-Patterson F_ST_ estimates between each pair of individuals within each species in R, to facilitate comparisons that do not rely on group assignments. With the group assignments and individual pairwise F_ST_ estimates, we statistically tested and graphically visualized relationships between these genetic measures and sampling year, sampling time of year, fish standard length, and juvenile status, to examine hypotheses about heterogeneity in spawning location and timing.

## Results

The GBS libraries yielded an average of 245 million reads across 200 individuals. The RAD libraries containing the *L. stappersii* individuals yielded an average of 271 million reads, including 2.5% – 8.6% bacteriophage PhiX genomic DNA. On average, the mapping rate for *L. stappersii* individuals’ GBS reads to the *L. calcalifer* reference genome was 89.8% (RAD, 89.6%), and mapping rates averaged 91.6% for *L. microlepis,* 91.3% for *L. mariae* and 88.7% for *L. angustifrons* individuals.

These mapping rates resulted in 1,463,702 unfiltered variable sites in *L. stappersii,* 1,286,810 in *L. microlepis,* 961,153 in *L. mariae* and 764,456 in *L. angustifrons.* After filtering for missing data (< 50%), minor allele frequency (MAF > 0.01), read depth (minDP > 5), and keeping only one SNP per locus (-thin 90), our species-specific GBS data sets contained 38,160 SNPs from 58 *L. stappersii* individuals (RAD, 16,802 SNPs from 63 different individuals), 33,429 SNPs from 34 *L. microlepis* samples, 30,431 SNPs from 38 *L. mariae* samples and 19,519 SNPs from 20 *L. angustifrons* samples. The data set with all four species contained 4,999,192 unfiltered and 44,823 filtered SNPs. None of the four species exhibited heterozygote excess that would be expected if many sites were erroneously called as heterozygous, and estimates of miscall rates are low for all read depths > 2 (Fig. S1). The majority of individuals who were lost were filtered out due to an imbalance in the reads supporting major and minor alleles at heterozygous loci. In our final dataset, the mean read depth was 15.2 for *L. stappersii,* 23.1 for *L. microlepis,* 20.9 for *L. mariae,* and 25.8 for *L. angustifrons* (Fig. S2).

### Species assignment

The species-combined SNP dataset for GBS data contained 200 individuals, of which we removed 50 individuals due to low numbers of reads (n=2), low assembly rate to the reference genome (n=4), or highly unbalanced read ratios at heterozygous sites (n=44), which is indicative of contamination. The remaining individuals all had > 50,000 reads that aligned to the *L. calcarifer* reference genome. The SNP alignment for RAxML contained 48,776 SNPs after removing invariant sites. The rooted maximum likelihood phylogeny (rooted on *L. calcarifer;* Fig. 2A) shows that each of the four species is reciprocally monophyletic. In addition, we infer a sister taxon relationship between *L. mariae* and *L. angustifrons,* and a sister taxon relationship between *L. stappersii* and *L. microlepis* (conflicting with the best supported topology in Koblmüller *et al*. 2021); however, short internode distances between these splits and low bootstrap support for the initial split suggest that all three speciation events may have occurred within a short period, and more investigation of the divergence history of the clade is warranted but not the focus of this current work. The PCA based on the GBS data was consistent with the species affinities indicated by monophyletic groups in RAxML (Fig. 2). The first genetic PC axis (27.3%) separates *L. stappersii* from the remaining three species. The second PC axis (12.4%) then separates *L. mariae* and *L. microlepis* from one another, and the third axis (4.8%) separates *L. angustifrons* from the other three species. Using entropy at K=4, we found each of the four species to make a distinct genetic cluster (Fig. 2C). Based on these combined analyses, 58 individuals were assigned to *L. stappersii,* 34 to *L. microlepis,* 38 to *L. mariae,* and 20 to *L. angustifrons.* While the genetic approaches revealed consistent species identifications, we had phenotypically misidentified one *L. microlepis,* one *L. mariae* and three *L. angustifrons* (Table S1).

**Figure 2.**
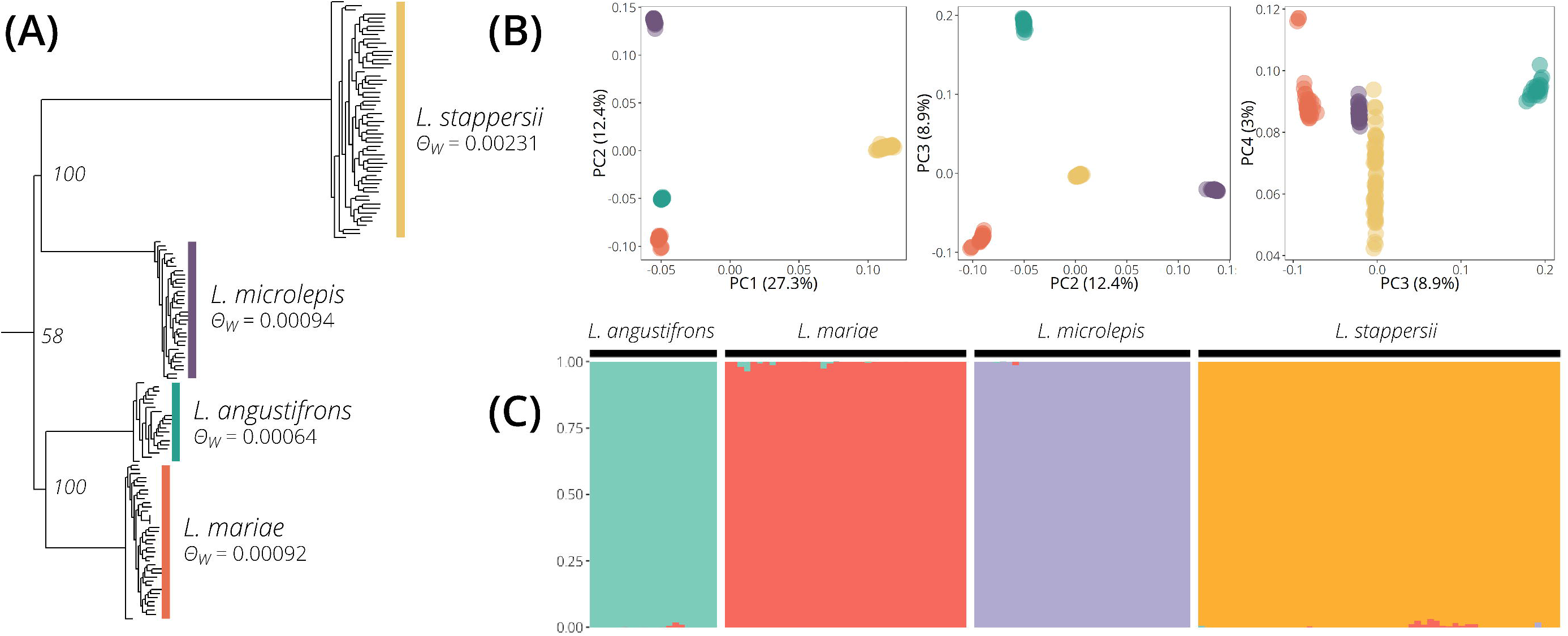
Relationships among the four endemic *Lates* species in Lake Tanganyika based on the GBS data set, as demonstrated by their (a) phylogenetic relationships, (b) relationships using principal component analysis, (c) genetic clustering using entropy at K=4. The phylogeny has been rooted using *L. calcarifer* as an outgroup and nodal bootstrap support is indicated for the main nodes separating the species. In (a), genetic diversity of the given species, as measured using Watterson’s Theta (*θ_w_*) in ANGSD, is indicated. Percentages in (b) indicate that amount of variance explained by the given principal component axis (axes 1-4 shown). In (c) each vertical bar represents an individual, and the colors represent the proportion of ancestry assigned to each of the four species. Individuals are colored by *a posteriori* genetic assignment of species, rather than phenotypic identification.

### Population structure

All four species had low but similar levels of genetic diversity (overall *θ_w_*= 0.00222), and genetic diversity was elevated in *L. stappersii* (*θ_w_*= 0.00230) compared to the three other species (*L. microlepis, θ_w_*= 0.00094; *L. mariae, θ_w_*= 0.00092; *L. anguistifrons, θ_w_=* 0.00064; Fig. 2A). Genetic diversity for the RAD dataset in *L. stappersii* was similar to that estimated for the GBS data (RAD *θ_w_*= 0.00223). Reflecting the distinctiveness seen in the all-species PCA and phylogeny, F_ST_ values are high between species (mean Reich-Patterson F_ST_ = 0.607; Table S3). None of the four species shows a significant relationship between genetic and geographic distance, suggesting an overall lack of isolation by distance (Fig. S3; all Mantel test *p*-values > 0.2).

The PCA for *L. stappersii* based on GBS data (Fig. 3A) did not reveal any clear genetic clusters. We similarly did not see any clustering in the RAD data for *L. stappersii* (Fig. S4). In entropy, K=1 was the best fit for the GBS data according to DIC (Fig. 5A); however, at K=2, some individuals from Kigoma, North Mahale and South Mahale form a distinct genetic cluster, and at K=3, groups appear that co-occur at all sampling sites. These groups at K=2 and K=3 were largely but not completely stable (i.e., the same individuals were clustered together) at a higher threshold for missing data (miss=0.5 vs. miss=0.9 had 18 individuals switching groups at K=2 and 10 individuals at K=3; Fig. S7, S8). This weak genetic structure (between-cluster K=2 Reich-Patterson F_ST_ = 0.00205, 95% bootstrap CI: 0.00084-0.00346, Table S4) was also partly reflected in F_ST_ values between sampling sites, where F_ST_ values of comparisons between individuals of South Mahale or North Mahale with samples from other sites were elevated, albeit small (≤ 0.012, Fig. 4A); however, only the F_ST_ values between North Mahale and each of Mpinbwe and Kagunga had 95% bootstrap confidence intervals not overlapping zero (Fig. S9A). The RAD data showed similar weak structure at K=2 (Fig. S5; between-cluster F_ST_ = 0.00146, 95% bootstrap CI: 0.00087-0.00207), but these groups were found in sympatry at all five sampling sites, similar to the groups at K=3 for the GBS data. At K=3, F_ST_ values between groups for the RAD data had bootstrap confidence intervals overlapping zero (Table S3).

**Figure 3.**
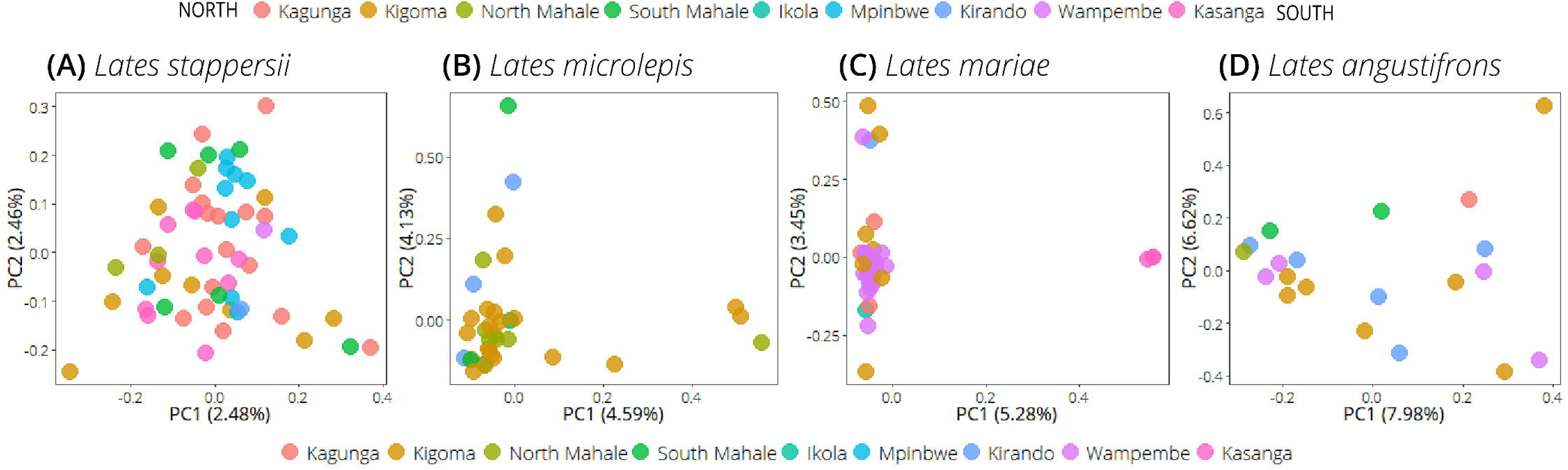
Principal component analysis of individual species based on the GBS data set, with colors indicating sampling locations, ordered from north to south. For each species, the plot shows the location of each individual on the two most dominant PCA axes, with percent variance explained in parentheses.

The PCA for *L. microlepis* did not reveal clear genetic clusters but it did highlight several individuals from the sampling sites Kigoma, North Mahale, and South Mahale, that were differentiated from the other samples on PC1 and PC2 (Fig. 3B). These differentiated individuals were not those with high amounts of missing data (Fig. S10B). All of the sampling sites had low differentiation (mean F_ST_ = 0.00471; Fig. 4B) but all pairwise comparisons had 95% confidence intervals that did not overlap zero except for the Kirando-Kigoma and Kirando-South Mahale comparisons (Fig. S9B). In entropy, K=1 was found to be the best fit using DIC (Fig. 5B); at K=2, four individuals collected in Kigoma are differentiated from the others, albeit with low genetic differentiation and a between-group F_ST_ estimate that overlaps zero (Reich-Patterson F_ST_ = 0.00074, 95% CI: −0.00389 – 0.00549; Table S4). At K=3, the additional group is found at all four sampling sites; the differentiation between the two largest groups is low, but significant (Reich-Patterson F_ST_ = 0.0030, 95% CI: 0.00155 – 0.00474), while the F_ST_ estimates between each of these groups and the smaller group identified at K=2 had 95% bootstrap confidence intervals overlapping zero. These groups at K=2 and K=3 were largely stable at a higher threshold for missing data (miss=0.5 vs. miss=0.9 had 4 individuals switching groups at K=2 and 8 individuals at K=3; Fig. S7, S8).

**Figure 4.**
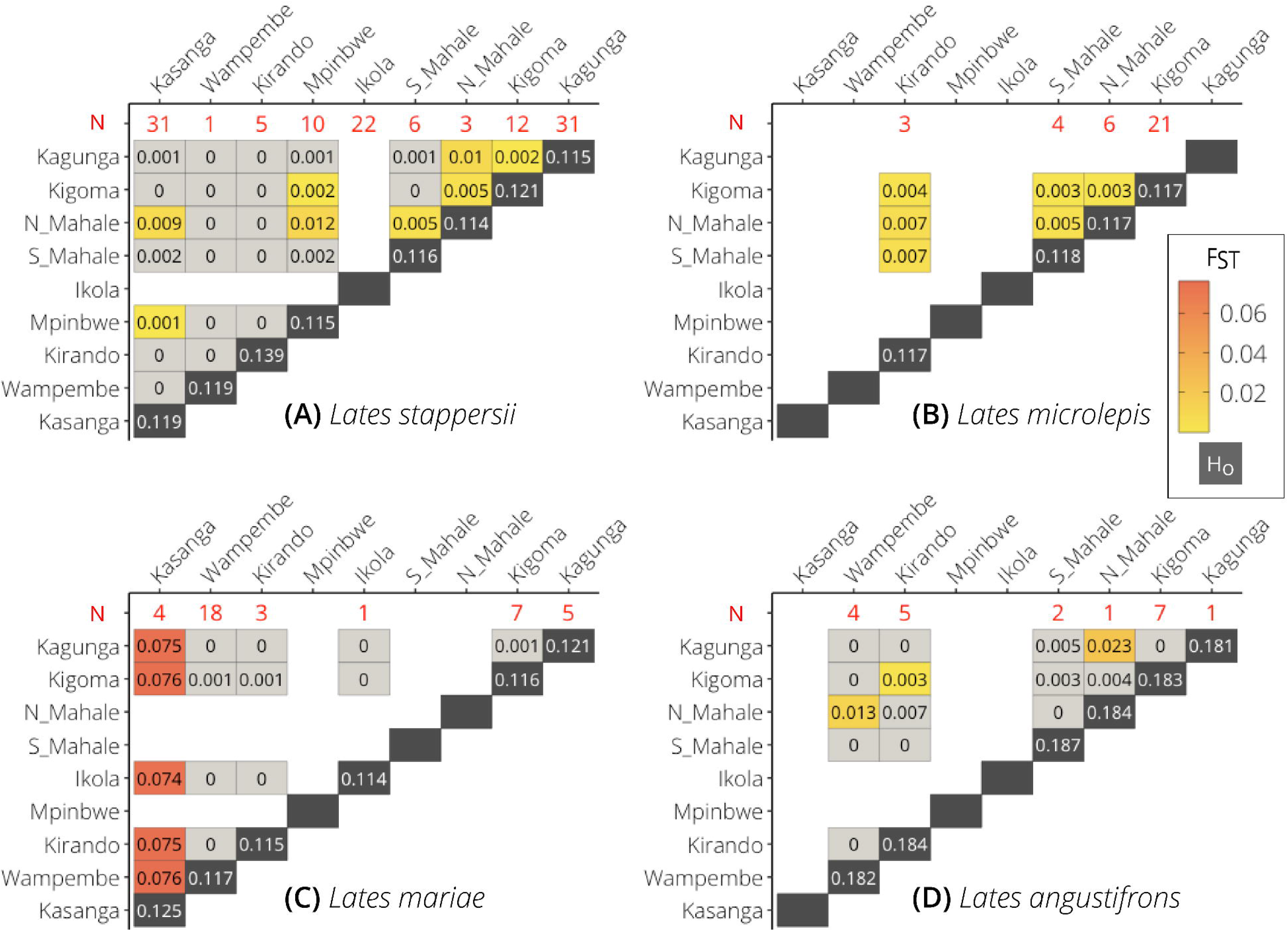
Heatmaps visualizing estimates of differentiation (Reich-Patterson F_ST_) between sampling locations above the diagonal, with observed heterozygosities (H_o_) along the diagonal (dark gray backgrounds) for the genotyping-by-sequencing dataset only. The red numbers across the top of each heatmap indicate sample size for the given species at the given sampling site. For F_ST_ values, light gray indicates no differentiation (i.e., bootstrap confidence intervals overlap 0), yellow indicates low differentiation, and red indicates relative high levels of differentiation.

**Figure 5.**
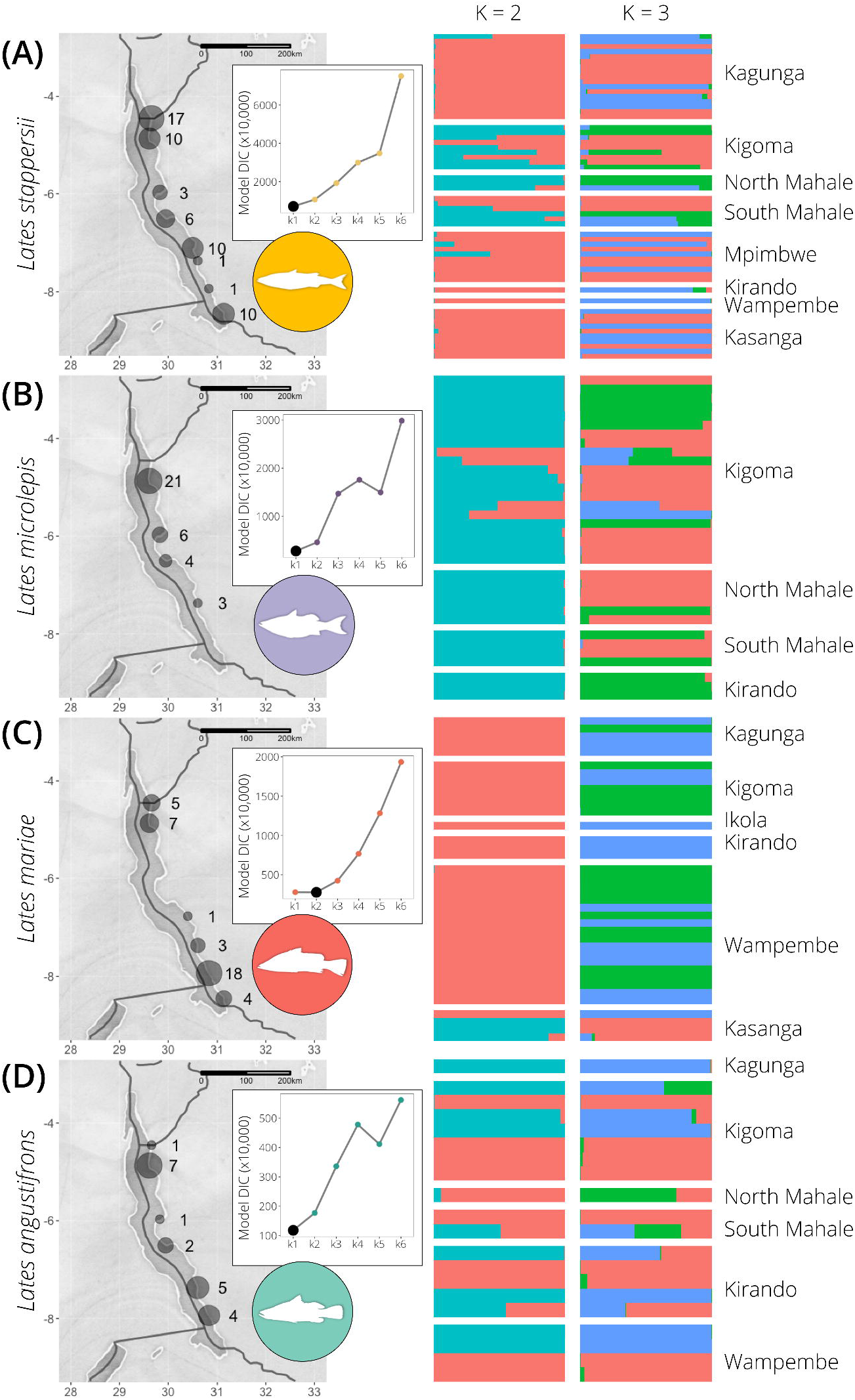
Intraspecific population structure inferred from the GBS data set and entropy analyses at K=2 and K=3 for (A) *L. stappersii,* (B) *L. microlepis,* (C) *L. mariae,* and (D) *L. angustifrons.* Maps show the distribution of individuals among sampling locations and inset plots show discriminant information content (DIC) values for each value of K, with the lowest DIC value for each species indicated with a larger black dot. In colored barplots, each bar corresponds to one individual and colors indicate inferred ancestry from each of the K groups. Plots for values of K=4 to K=6 can be found in Supplementary Fig. S6.

The PCA for *L. mariae* clusters three individuals from Kasanga separate from all other individuals on the first PC axis (explaining 5.28% of the variance), while the second PC axis separates out some individuals from Kigoma, Wampembe, and Kirando from the others (Fig. 3C). The differentiation among individuals on both PC1 and PC2 does not correlate with the individuals’ amount of missing data (Fig. S10C). In entropy, K=2 has the lowest DIC estimate, where the three individuals from Kasanga form a distinct cluster (Reich-Patterson F_ST_ between entropy-identified groups = 0.119, 95% bootstrap CI: 0.113-0.127; Fig 4C and Table S4). At K=3, most of the other sampling sites have individuals assigned to both of the non-Kasanga clusters, although these two non-Kasanga clusters have very low differentiation (F_ST_ = 0.00359, 95% bootstrap CI: 0.00180-0.00623) compared to that between the Kasanga cluster and each of the other two clusters (mean F_ST_ = 0.120). These groups at K=2 and K=3 were largely stable at a higher threshold for missing data (miss=0.5 vs. miss=0.9 had no individuals switching groups at K=2 and 15 individuals at K=3; Fig. S7, S8). The F_ST_ estimates between all fish sampled from Kasanga and other sampling sites are all significantly different from zero (Fig. S9C). All other between-site F_ST_ estimates had 95% bootstrap confidence intervals overlapping zero.

For *L. angustifrons,* the PCA did not reveal any genetic structure (Fig. 3D) and the clustering model with K=1 is the best fit for our data (Fig. 5D). Clusters identified at higher levels of K had low differentiation (Table S4; K=2, F_ST_ = 0.00498; K=3, mean F_ST_ = 0.0228), and were found co-occurring at most sampling sites (Fig. 5D). These groups at K=2 and K=3 were largely but not completely stable at a higher threshold for missing data (miss=0.5 vs. miss=0.9 had 4 individuals switching groups at K=2 and 7 individuals at K=3; Fig. S7, S8). In addition, only three pairwise F_ST_ estimates between sampling sites were significant: Kagunga-North Mahale, Kigoma-Kirando, and North Mahale-Wampembe (Fig. S9D); however, even these sites had low differentiation (mean F_ST_ between sites = 0.0132), and it should be noted that North Mahale only has one fish sampled, so these estimates may not be indicative of patterns at the site as a whole.

We find no relationship between size difference and extent of differentiation (pairwise F_ST_ or genetic cluster membership) in *L. mariae, L. microlepis,* or *L. angustifrons.* In *L. stappersii,* a weak positive relationship exists between the difference in standard length and the extent of genetic differentiation (F_ST_) between individuals, such that individuals that are more different in size are also more differentiated genetically (Fig. S12). In *L. microlepis,* we found no relationship between genetic cluster membership and juvenile status at K=2 or K=3 when examining only the fish collected in Kigoma, the only site with large enough numbers for doing this analysis. We additionally find no systematic difference between F_ST_ estimates between only juvenile fish and those estimated between only adult fish for *L. microlepis* or *L. mariae* (Fig. S13, S14). When only looking at fish collected at the same site (Fig. S14), we find that juvenile-adult comparisons had higher F_ST_ values in *L. mariae* than adult-adult comparisons (two-tailed t-test: t = 2.3484, df = 28, p-value = 0.02615), but did not differ from juvenile-juvenile comparisons (two-tailed t-test: t = −1.7863, df = 31.781, p-value = 0.0836), and juvenile-juvenile and adult-adult comparisons were not significantly different from one another (two-tailed t-test: t = 1.9658, df = 151, p-value = 0.05116). In *L. angustifrons,* F_ST_ values among juveniles at the same site were lower than those between juveniles and adults at the same site (two-tailed t-test: t = −3.8334, df = 16.646, p-value = 0.001376) or between adults at the same site (two-tailed t-test: t = 2.2254, df = 27.295, p-value = 0.0345), while F_ST_ estimates did not differ significantly between juvenile-juvenile and adult-adult comparisons (two-tailed t-test: t = −1.6493, df = 14.084, p-value = 0.1212). In *L. microlepis,* F_ST_ estimates for fish from the same site were marginally higher in juvenile-juvenile pairs than in adult-adult pairs (two-tailed t-test: t = 2.2525, df = 19.005, p-value = 0.03631), but did not differ between each of these categories and juvenile-adult comparisons (p-value > 0.07). We do not find any evidence for juveniles collected at the same time of year to be more closely related than juveniles collected during different parts of the year in *L. microlepis* (two-tailed t-test: t = – 1.2229, df = 378.64, p-value = 0.2221; Fig. S15), and do not have enough data to analyze this relationship in the other three species.

## Discussion

Differences in life history characteristics can have significant influence on variation in population genetic structure among taxa, even in closely related species. In this study, we analyzed population genetic structure within the four endemic pelagic top predators of Lake Tanganyika, *Lates spp.,* using reduced-representation genomic sequencing data sets. From our samples collected at discrete sampling sites along the length of the lake, we find two key patterns: first, we find a strongly differentiated genetic group in *Lates mariae* at the most southern sampling site, Kasanga; second, we find no additional pattern of isolation-by-distance or geographic population structure within any of the four *Lates* species. We find weak evidence for weakly differentiated genetic groups that exist together at the majority of sites in all four species. While numerous explanations could account for a lack of geographic population structure and the existence of sympatric weakly differentiated groups, more research is needed to understand mechanisms causing the population structure identified in these species. Our results have potential implications for understanding the conservation and management of these fishes, with implications for this regionally important fishery.

### Strong genetic differentiation in southern Lates mariae

In *L. mariae,* the majority of samples from the most southern site, Kasanga, are strongly genetically differentiated from all other samples. The magnitude of F_ST_ estimates between these distinct individuals and the other *L. mariae* individuals (F_ST_ ~0.135), suggests unexpectedly strong intraspecific genetic structure given a lack of obvious geographic barriers. Our sampling does not include *L.* microlepis or *L. angustifrons* individuals from this furthest south site, so we cannot use our current data to determine whether this pattern is also reflected in the other two large *Lates* species.

It is plausible that the differentiation in *L. mariae* may be driven by limnological patterns that differ between the northern and southern portions of the lake. In particular, the southern part of the lake experiences intense upwelling of cold, nutrient rich waters from the deep during the dry season (Verburg *et al*. 2011), which mixes with oxygenated water from the surface and consequently has high nutrient concentrations and primary productivity rates, which in turn fuels zooplankton growth (Plisnier *et al*. 2009; Loiselle *et al*. 2014). In contrast, the northern part of the lake has a more stable, persistently stratified, clear water column (Plisnier *et al*. 1999; Bergamino *et al*. 2010; Mziray *et al*. 2018). As a result, there is asynchronous variability between environments in the north and south, as well as high environmental variability over the course of the year in the south (i.e, higher primary productivity in the dry season compared to the rainy season). This spatial variability affects zooplankton community composition, with shrimps and calanoid copepods prevailing in the south (Kurki *et al*. 1999), while cyclopoid copepods and medusa dominate in the north (Kurki *et al*. 1999; Cirhuza and Plisnier 2016). Increases in primary productivity generally lead to high densities of zooplankton, as well as *Stolothrissa* and *Limnothrissa* sardines (Mulimbwa *et al*. 2014), as the blooms seem to positively affect sardine spawning and recruitment, although recruitment pulses sometimes occur during times of poor nutrient conditions. Mannini *et al*. (1999) found corresponding spatial variability in diet for *L. stappersii,* but spatial variability in diet has not been documented for *L. mariae* or the other large-bodied species. While adult *L. mariae* do not depend on the pelagic sardines as a primary food source (Coulter 1976), it is possible that such temporal nutrient variability leads to variability in the main *L. mariae* prey (benthic cichlids) as well.

Such a spatially and temporally variable environment might lead to local differential selection in the southern part of the lake, and this selection combined with restricted movement of individuals between this and other populations of *L. mariae* could lead to the genetic differentiation in individuals from Kasanga. *L. mariae* abundance is greater in the south than in the north (Coulter 1970; personal communication), which suggests that spatial variation in environmental conditions may underlie differences in fitness or population growth. A variable environment may also lead to within-site differentiation among individuals caught in Kasanga if the two groups were to adapt to this variability in different ways.

Another possibility is that one group is resident at Kasanga, while the other disperses throughout the lake. While all *L. mariae* individuals sampled from Kasanga were mature and *L. mariae* is believed to be the least vagile of the four *Lates* species (Coulter 1976), it is not clear how vagile these individuals typically are; it is possible that some individuals remain resident while others migrate, as is the case in Atlantic cod (Kirubakaran *et al*. 2016; Berg et al. 2017). To investigate these possibilities, future studies should expand sampling in the southern portion of the lake, including sampling various life stages of this and other *Lates* to understand the genetic composition of the southern populations and any important ecological implications of genetic differentiation.

It is also possible that the differentiation in the most southern part of Lake Tanganyika in *L. mariae* is related to the lake’s bathymetry and differences in the depth of the oxygenated layer, as suggested by Coulter (1976), who observed asynchronous changes in local abundance between *L. mariae* populations in the southwest versus southeast arms of the lake. From catch rates, Coulter (1970) observed that southern populations of at least two of the *Lates* species (*L. angusitfrons* and *L. mariae)* were likely distinct, as they were heavily exploited in the southeast arm with no apparent effect on catches in the southwest arm. These southern reaches of the lake are relatively shallow, and the oxygenated layer extends up to 200m deep, which allows for an abundance of oxygenated habitat for benthic species (Coulter 1976).

Despite the life history differences between pelagic cichlids and the *Lates* species, the genetic patterns observed in *L. mariae* are similar to those observed previously in the benthopelagic cichlids. For the four benthopelagic *Diplotaxodon* species of Lake Malawi with genetically structured populations, research by Genner *et al*. (2010) indicated spawning site fidelity drives population structure. In contrast, Koblmüller *et al*. (2019) hypothesized that preying upon benthic cichlids—which requires shorter-distance movements than preying on pelagic prey—can explain the genetic structure in the *Bathybatini* of Lake Tanganyika. Similar to the *Bathybatini, L. mariae* hunts benthic cichlids and is described as relatively resident (Coulter 1976). However, our results did not recover any evidence for significant isolation-by-distance in *L. mariae,* suggesting that the genetic structure that we observe is likely not the result of dispersal limitation. Interestingly, females of *L. mariae* likely aggregate over specific spawning grounds (Coulter 1991), similar to *Diplotaxon.* If samples of *L. mariae* were taken from spawning grounds, it would be possible to infer whether the divergence we observe is related to strong spawning site fidelity and differentiation between these stocks. However, the sampling methods used in our study (i.e., obtaining fish from fishermen) limit our knowledge of exact catch location. In addition, these sampling methods mean that a range of ages and reproductive maturities of fish were included in our study. To our knowledge, the location of spawning grounds has not been documented for any of the four *Lates* species. In addition, it is unclear how the pelagic phase of eggs, larvae, and small juveniles (<2.5cm), of *L. mariae* influences dispersal in the species. Spawning pelagically (Riginos *et al*. 2014) and having a long pelagic larval stage (Selkoe & Toonen 2011; Shanks 2009) generally facilitate gene flow and reduce population structure in marine taxa, although these patterns are complicated by specific behavior exhibited by both adults and larvae (Bay *et al*. 2006; Riginos *et al*. 2011). Future studies in *L. mariae* should sample spawning fish and work to understand how genetic divergence relates to spawning site location and timing.

### No pattern of isolation-by-distance or geographic population structure

Although the samples spanned the ~550 km Tanzanian shoreline of Lake Tanganyika, none of the genetic differences between populations relates to geographic distance, suggesting no isolation-by-distance in any of the four species. In addition, admixture models with only a single panmictic population fit our data the best in all species except for *L. mariae* (Fig. 5). This suggests that individuals of all four species are mobile enough that gene flow is not restricted among our sampling sites, other than the distinct Kasanga population.

Deviations from isolation-by-distance are common among marine animals, where geographically proximate populations often have greater population differentiation than populations further apart, a phenomenon sometimes referred to as “chaotic patchiness” (Johnson and Black 1984). This phenomenon has been attributed to larval dispersal via ocean currents (White et al. 2010), and it is possible that currents within Lake Tanganyika may similarly disperse *Lates* eggs and larvae prior to their movement to inshore habitats. While it is presumed that all four *Lates* species spawn pelagically (Ellis 1978; Coulter 1991) and the larvae of the larger three species are planktonic until ~3cm in size (Coulter 1976), spawning behavior, spawning location within the lake, and the length of time that larvae may be carried around by lake currents remain undocumented for all four species. Lake currents vary throughout the year (Plisnier *et al*. 1999; Verburg *et al*. 2011), and therefore understanding the timing and location of reproduction for all four species will be important to understanding the role that these currents may play in dispersing planktonic eggs and larvae.

While geographically widespread populations are not particularly surprising given the large body sizes and long-distance swimming capabilities of these fishes, it conflicts with evidence of a resident lifestyle in *L. mariae* (Chapman 1976; Coulter 1976, 1991; Muller *et al*. 2001) and evidence for some spatial structure in the Kigoma area in a previous population genetic study of *L. stappersii* (Kuusipalo 1999). It should be noted that our sampling is unbalanced among sites and lacking in the most southern reaches of the lake for *L. angustifrons* and *L. microlepis.* It is therefore possible that more (albeit still likely weak) genetic structure exists than we have been able to detect with available samples. That said, Koblmüller *et al*. (2019) similarly found no evidence of genetic population structure in two eupelagic bathybatine cichlid species of Lake Tanganyika, Koblmüller *et al*. (2015) found no evidence for population structure in the pelagic cichlid *Boulengerochromis microlepis,* and De Keyzer *et al*. (2019) and Junker *et al*. (2020) found no population structure in either of the pelagic sardine species. There are many life history differences between the sardines, cichlids, and *Lates* spp. that occur sympatrically in the open waters of Lake Tanganyika, but these results together suggest that spatial environmental heterogeneity along the length of the lake is not sufficient to restrict gene flow and produce geographically isolated or clinal populations in these diverse species.

### Sympatric, weakly-differentiated groups exist in all four species

While our admixture models suggest panmixia as the best model in *L. stappersii, L. microlepis,* and *L. angustifrons,* and two distinct groups in *L. mariae,* we find sympatric differentiation in all four *Lates* species at higher K values. Model DIC values suggest that these higher K values are not the best fit for our data, and differentiation is weak (mean F_ST_ = 0.032; Table S4), but individual assignments do not behave as would be expected for truly panmictic populations. In our data, we see individuals with high assignment confidence to one group or another (i.e., each bar is mostly a single color in Fig. 5 and Fig. S6); if the population were truly panmictic, then we would expect to see all individuals showing genetic contributions from all groups at higher than optimal values of K. For that reason, we believe these weakly differentiated groups may be biologically relevant, albeit weakly differentiated. These groups do not appear to correspond to sex or any other phenotypic character that we recorded. There are several potential explanations for this consistent structure, of which we will discuss the evidence for three: (a) spatially segregated spawning sites with spawning site-based differentiation, where adults move throughout the lake but return to the same spawning grounds to breed; (b) temporally segregated spawning groups; or (c) historical allopatric groups that have become sympatric yet partially retained their genetic distinctiveness.

Genetically distinct groups that occur in sympatry must have at least partial reproductive isolation, and in fish without clear sexual selection, this often occurs intraspecifically as a segregation of spawning either in space or time. If there is site fidelity to regionally differentiated spawning grounds but between-region movement outside of spawning, then this could create patterns of differentiation such as those that we observed. If spatially segregated spawning sites exist, then the adults that are sampled at a single site may have migrated from a more distant site (unless they are collected while spawning, in which case we would expect the genetics to match with their geographic location), but we would expect littoral juveniles to exhibit a stronger pattern of geographic-based differentiation or isolation by distance. In addition, we would expect individual differentiation among juveniles collected at the same site to be lower than differentiation among adults collected at the same site, or differentiation between juveniles and adults collected at the same site. We did not find this to be the case in any of the four species (Fig. S14), with the caveat that our sample sizes for these comparisons are low and our data may exhibit batch effects due to our sampling strategies. We did find stronger differentiation in *L. angustifrons* and *L. mariae* between juveniles and adults collected at the same site than among juveniles collected at the same site; however, the mean F_ST_ for adult-adult comparisons was not larger than that of juvenile-juvenile comparisons, so the evidence supporting this hypothesis is mixed.

Genetic differentiation related to temporal stratification in spawning behavior—such as having a population of “dry season” spawners and a population of “rainy season” spawners— could also explain the existence of sympatric groups. Coulter (1976) suggested that the four species likely spawn continuously, but with peaks in number of ripe females from August to November/December, and potentially other smaller peaks in March/April. In addition, observations of well-differentiated size distributions of young *Lates* found in littoral weed beds suggests periodicity in spawning (Coulter 1976). Unfortunately, we do not have data on the spawning status of all of the fish collected in this study, which would facilitate investigating this hypothesis. However, we were able to test whether juveniles collected at a similar time of year were more closely related than those collected at a different time of year, regardless of sampling site. We do not find any evidence for juveniles collected at the same time of year to be more closely related than juveniles collected during different parts of the year in *L. microlepis* (Fig. S15); however, our sampling is not conducive to being able to rigorously investigate this relationship, as some sites were only sampled in one year or at one time of year. Additionally, only a few juveniles were collected opportunistically from each species, as they are too small to be targeted by fishermen. Investigating the possibility of discrete spawning groups will be an important next step in understanding the weak genetic structure present in these species, and will require more targeted sampling.

A third potential explanation for the intraspecific structure could lie in the geologic history of Lake Tanganyika. With an estimated age of 9-12 million years, Lake Tanganyika is one of the oldest lakes in the world and has been influenced by extensive lake level fluctuations (Cohen *et al*. 1993). Based on the bathymetry of the lake, it is thought that Lake Tanganyika was divided into three paleo-lakes during periods of extremely low water levels (Cohen *et al*. 1997). Such environmental fluctuations may have led to oscillations between sympatry or parapatry and allopatry for fish populations within the lake, which have been invoked as a “species pump” (Greenwood 1981; Sedano *et al*. 2010) contributing to diversity in some clades in the littoral (Janzen & Etienne 2017; Sturmbauer *et al*. 2016) and benthic or pelagic (Duftner *et al*. 2005; Koblmüller *et al*. 2005) zones of the lake. While it has recently been suggested that the origin of the Lake Tanganyika *Lates* radiation is much younger than other endemic fish radiations in the lake (Koblmüller *et al*. 2021), it remains unclear whether speciation in this clade was similarly triggered by these environmental fluctuations—as well as whether more recent fluctuations could explain our contemporary observations of weak genetic structure. Such fluctuations could also lead to persistent genetic structure within species, for example, if spawning site fidelity within separate basins is maintained after the basins reconnect. However, we would expect strong philopatry to produce much stronger genetic differentiation than observed here.

Interestingly, *L. stappersii* in this study which were caught during the dry season in the south of the lake show distinct carbon isotope signals from samples caught in the north (Ehrenfels *et al*. 2021). These differences in carbon isotope ratios correlate with different rates of primary productivity in the northern and the southern basin during the dry season (Loiselle *et al*. 2014), and with differences in prey composition in different areas of the lake (Plisner *et al*. 2009; Kurki *et al*. 1999). This is evidence for at least partly resident populations of *L. stappersii,* which stay in the northern or southern basin during the dry season for long enough to incorporate the different isotopic signals of these basins. Such movement patterns may reduce gene flow between *L. stappersii* populations from the basins; however, these isotopic groups do not correspond with the genetic groups found at K=2 here (Ehrenfels *et al,* 2021).

It is also important to keep the limitations of these data in mind. Admixture analyses using programs such as STRUCTURE (Pritchard *et al*. 2000) are biased in cases of unbalanced sampling among sites, such that differentiation is often overestimated at sites with the most individuals and underestimated at sites with few individuals (e.g., Puechmaille 2016; Wang 2017). While it is unclear whether this is also true for entropy analyses, it is possible that a similar issue influences our results. For example, we find multiple groups at the Kigoma site in *L. microlepis* at K=2, which is also the site with the most individuals (Fig. 5B). Furthermore, our assignment of individual locations to fisherman landing sites or markets is less precise than if we had exact locations where the fish were caught. While we only have general knowledge of how far fishermen travel from the landing site to fish during the day, it is possible that fish collected at the same sampling site were collected up to ~40 km away from one another. In addition, our samples collected in the Kigoma fish markets may have come from farther away from those collected at landing sites (Petit & Shipton 2012). These limitations may influence our ability to detect geographic trends in our data.

### *Recommendations for conservation and sustainable management of Lake Tanganika’s* Lates

We find evidence for weak differentiation within sampling sites in all four species. We furthermore find clear evidence for a genetically distinct group within *L. mariae* at our most southern sampling site, in agreement with previous studies suggesting that this species is the most resident of all four *Lates* species (Coulter 1976, 1991). Intriguingly, the weak intraspecific structure in all four species is not linked to isolation-by-distance, but rather involves differentiation between some individuals at multiple sites, suggesting the existence of a mechanism for maintaining differentiation while these distinct genetic groups are in sympatry. In contrast, the low F_ST_ estimates between samples from distant sites and lack of isolation-by-distance suggest that these cies disperse over large distances, indicating a need for management of these species on at a lake-wide scale. Furthermore, it is likely that more extensive sampling of these little-studied species may uncover important additional diversity.

To maintain intraspecific genetic variation, viable population sizes, and possible adaptive differences between genetic groups within the *Lates* species, more research is needed on the location and timing of spawning for the four species and whether restricting catches in certain areas and periods of the year could help the larger-bodied species recover from low population sizes. Harvest restrictions have been implemented in regions around Lake Tanganyika where community conservation areas are active (Kimirei & Sweke 2018) but are not in place throughout the lake. Such management practices are performed successfully in the cod fishery of Denmark (albeit after collapse of this fishery and extinction of many of the original stocks; Dahle *et al*. 2018), and in the salmon fishery of Bristol Bay (Schindler *et al*. 2015; Schindler *et al*. 2010). However, implementation of such a strategy requires detailed evidence of spawning ground locations and timing of spawning, as well as for fishermen to be able to distinguish between the genetically distinct stocks. Therefore, studies are needed to shed more light on the phenotypic, ecological, and phenological characteristics of the genetic groups within each species. In addition, studies are needed to examine whether fish from different spawning grounds form the distinct genetic clusters that we identified in this study. Genomic analyses, although helpful in generating hypotheses about the natural history of these species, must be paired with an in-depth understanding of natural history, including integration of local knowledge, for managing these fish stocks. A sustainable management approach informed by these combined data is crucial to protect the inshore nursery habitats and ensure recruitment of the species in Lake Tanganyika.

## Funding

This work was supported by the Swiss National Science Foundation (grant CR23I2-166589 to OS), the University of Wyoming (start-up funding to CEW), the National Science Foundation (grant DEB-1556963 to CEW), the National Institute of General Medical Sciences of the National Institutes of Health (Institutional Development Award grant P20GM103432), and the American Genetics Association (Ecological, Evolutionary, and Conservation Genomics Research Award to JAR).

## Acknowledgements

Computing was accomplished with an allocation from the University of Wyoming’s Advanced Research Computing Center, on its Teton Intel x86_64 cluster (https://doi.org/10.15786/M2FY47) and the Genetic Diversity Center (GDC) of ETH Zürich. Special thanks go to Benedikt Ehrenfels for coordinating field campaigns, Mupape Mukuli for facilitating logistics during fieldwork, and to the crew of the MV Maman Benita for fieldwork assistance. We thank the whole team at the Tanzanian Fisheries Research Institute for their support. A special thanks goes to Mary Kishe for her support during fieldwork permission processes and to the Tanzanian Commission for Science and Technology (COSTECH) for the permits to do this research. Live fish samples were collected under the approved University of Wyoming IACUC protocol #20160602CW00241-01. We appreciate The Nature Conservancy’s Tuungane Project for providing funding and logistical support, with special thanks going to Colin Apse and Peter Limbu. We also thank the members of the Wagner Lab at the University of Wyoming and the FishEc group at EAWAG for discussions about the analyses and manuscript. We also thank the Associate Editor and four anonymous reviewers for the valuable comments that greatly improved the quality of this manuscript.

## Data Availability

We have deposited the primary data underlying these analyses as follows:

- Sampling locations, morphological data, and filtered VCF files have been archived on Zenodo (http://doi.org/10.5281/zenodo.5216259)
- Fastq files for each individual have been archived in NCBI SRA PRJNA776855, and will be released upon publication
- Scripts for data analysis are available on Github (https://github.com/jessicarick/lates-popgen), and the version at publication will be archived on Zenodo (http://doi.org/10.5281/zenodo.5216259)

## Notes

http://doi.org/10.5281/zenodo.5216259

### Competing Interest Statement

The authors have declared no competing interest.

### Summary of Updates

Minor changes to text to improve readability.

