## Supplemental Tables and Figures for "The genetic population structure of Lake Tanganyika’s *Lates* species flock, an endemic radiation of pelagic top predators"

**Table S1.** Species identification based on phenotype and the genetic assignment methods. In *pheno_ID*, species identification is indicated from following FAO phenotypic characters ("juvenile" indicates that the fish is too young to distinguish species based on these characters). "PCA_ID_GBS" and "PCA_ID_RAD" indicate species identity from clustering in principal component analysis; entropy_ID indicates species identified from admixture analysis in entropy; "final_ID" indicates species designation from combining all lines of evidence. The "checkHets_qual" column indicates the ratio of reads supporting the minor versus major allele at heterozygous sites (from 1=best to 4=worst). For phenotypic characters, "SL_mm" is standard length in millimeters, "TL_mm" is total length in millimeters, and "mass_g" is mass in grams.

*See separate Rick_SupMat_TableS1.csv file for Table S1*

**Table S2.** Estimates of $F_{ST}$ and 95% confidence intervals between species pairs, calculated using the Reich-Patterson $F_{ST}$ estimator. Confidence intervals come from 100 bootstrap replicates.

| | *L. stappersii* | *L. microlepis* | *L. mariae* | *L. angustifrons* |
|---|---|---|---|---|
| *L. stappersii* | | **0.691**<br>95% CI:<br>0.686-0.696 | **0.668**<br>95% CI:<br>0.662-0.673 | **0.693**<br>95% CI:<br>0.687-0.698 |
| *L. microlepis* | | | **0.651**<br>95% CI:<br>0.646-0.656 | **0.681**<br>95% CI:<br>0.673-0.688 |
| *L. mariae* | | | | **0.603**<br>95% CI:<br>0.595-0.611 |
| *L. angustifrons* | | | | |

**Table S3**. Mean Reich-Patterson $F_{ST}$ among intraspecific groups identified in entropy analyses for each species from K=2 to K=6. Bolded numbers indicate that all 95% confidence intervals for $F_{ST}$ estimates did not overlap zero.

| | **K = 1** | **K = 2** | **K = 3** | **K = 4** | **K=5** | **K=6** |
|---|---|---|---|---|---|---|
| *L. stappersii* | | **0.00409**<br>95% CI: 0.00284-0.00517 | **0.00542** | 0.00326 | 0.00434 | 0.00384 |
| *L. microlepis* | | 0.00074<br>95% CI: -0.00343 - 0.00456 | 0.00241 | 0.00274 | 0.00232 | 0.00314 |
| *L. mariae* | | **0.135**<br>95%CI: 0.128-0.140 | **0.0927** | 0.0702 | 0.0717 | 0.0563 |
| *L. angustifrons* | | **0.0325**<br>95% CI: 0.0295-0.0352 | 0.0228 | 0.0193 | 0.0133 | 0.0237 |

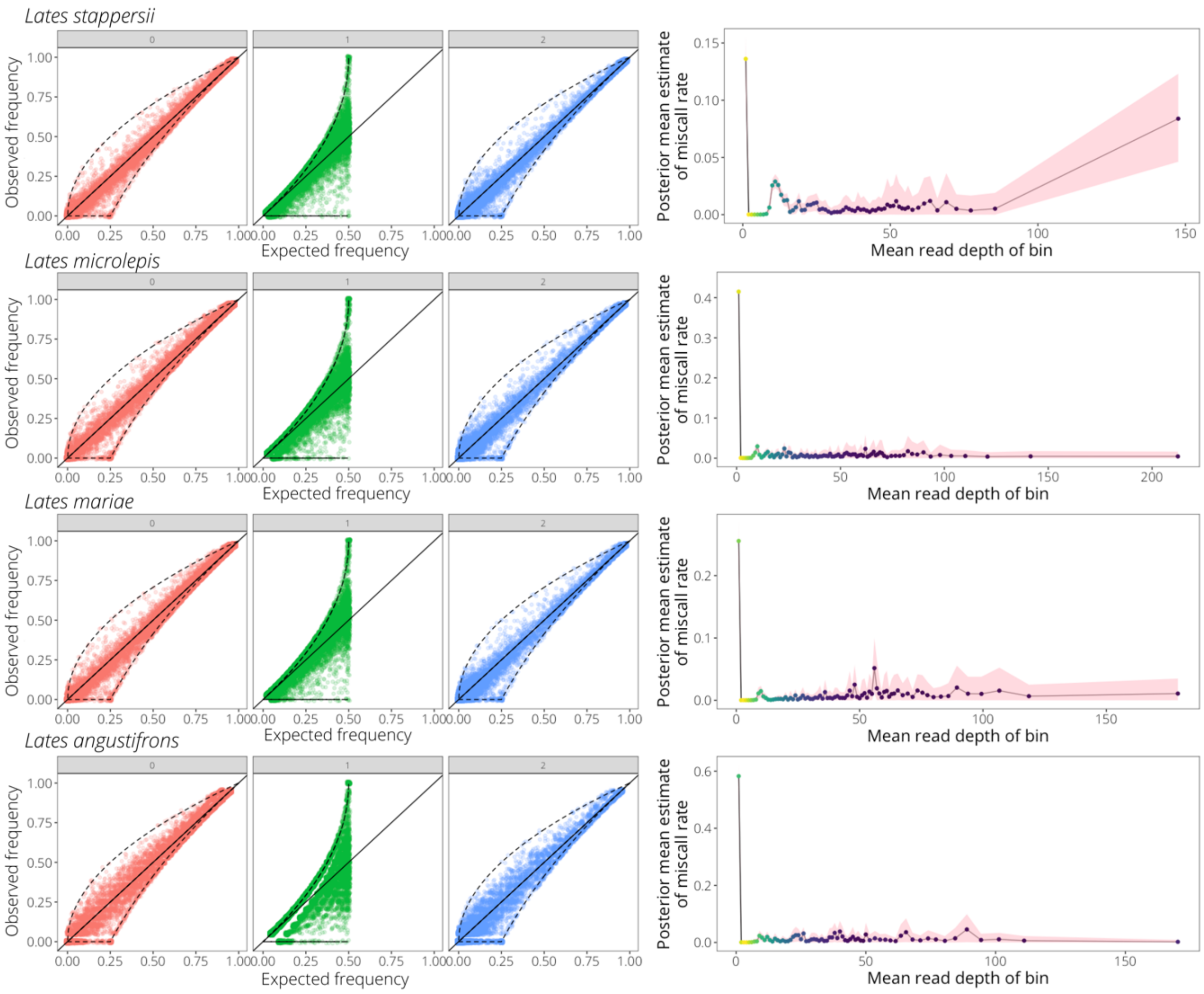


**Figure S1**. Assessment of heterozygote excess and modeled estimates of heterozygote miscall rate at increasing mean read depths for each of the four species, as assessed using whoa (Anderson 2018) prior to filtering for minimum read depth. Loci largely conform to Hardy-Weinberg expectations for homozygotes and heterozygotes, and miscall rates are only large in the smallest read depth bin (0 < mean read depth < 1).

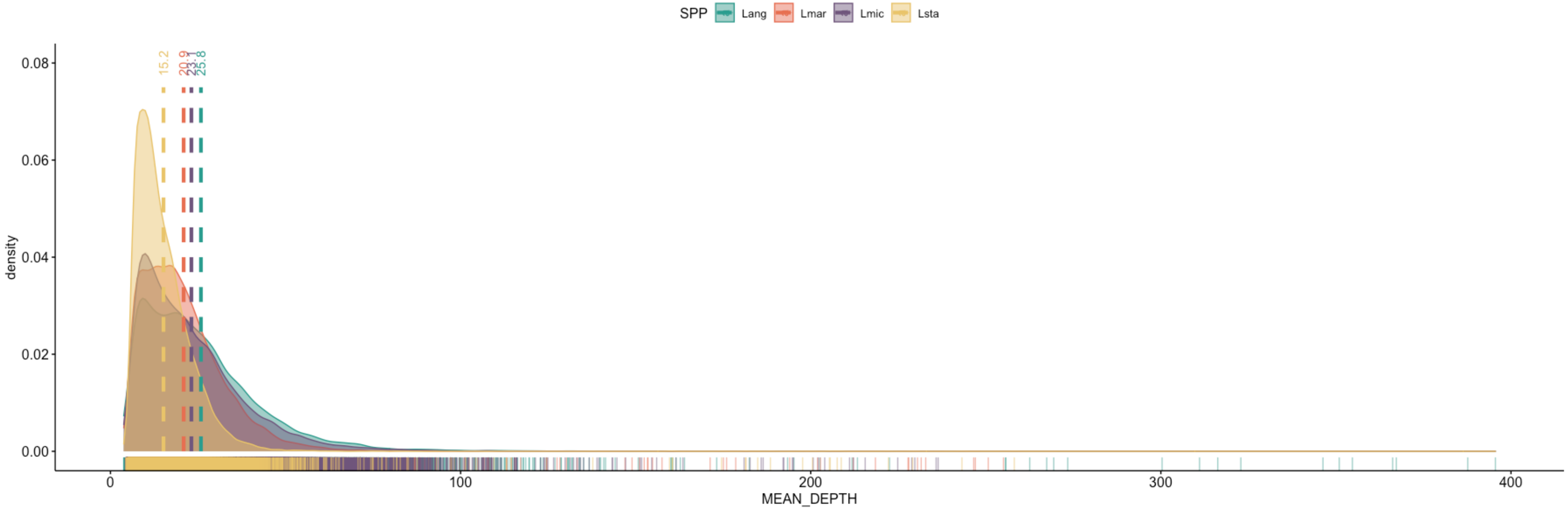


**Figure S2**. Post-filtering distribution of read depths for each species, with mean read depth across sites indicated by vertical dashed lines (*L. stappersii*, mean=15.2; *L. microlepis*, mean=23.1; *L. mariae*, mean=20.9; *L. angustifrons*, mean=25.8)

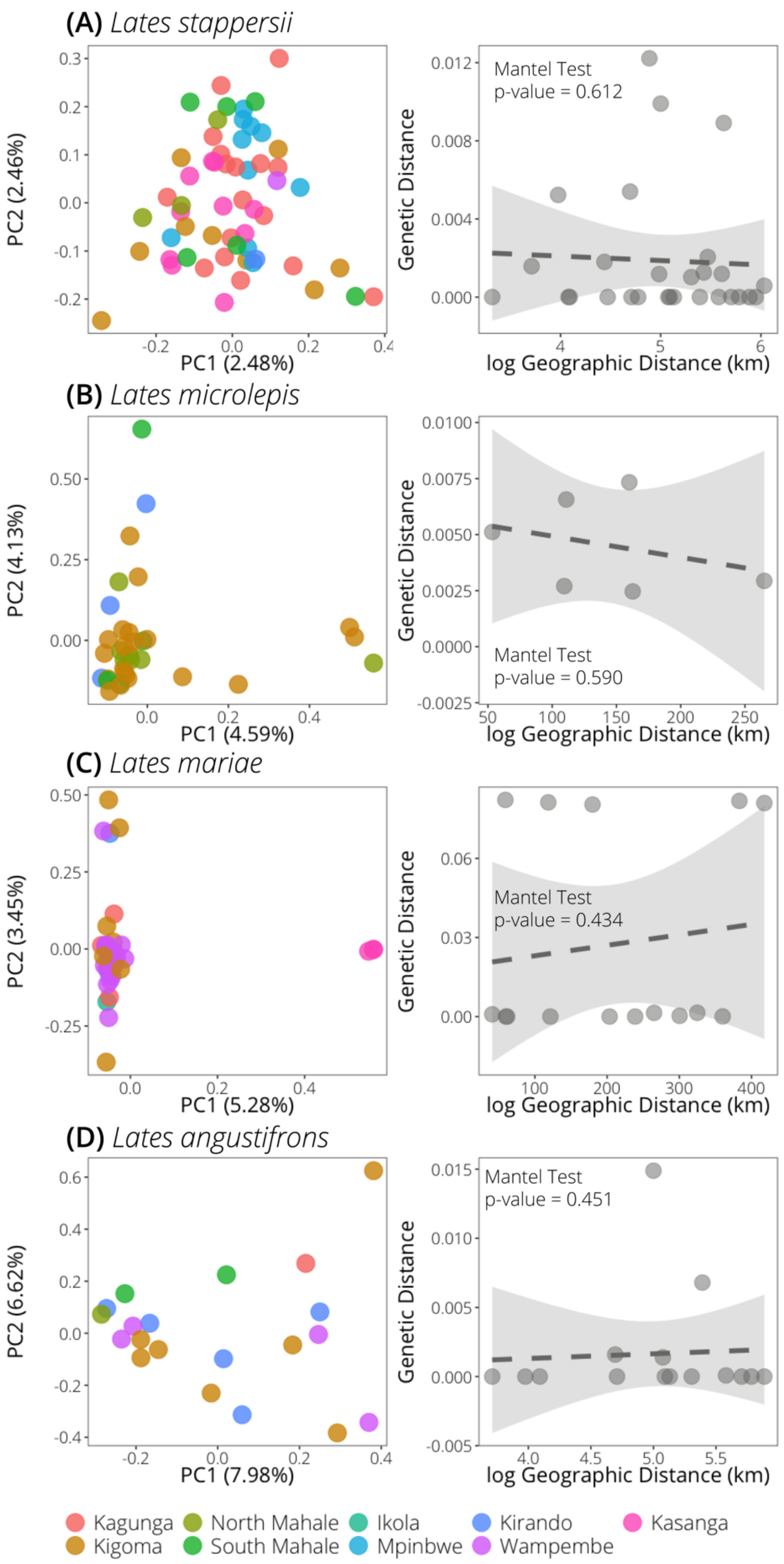


**Figure S3.** Principal component analysis and isolation-by-distance plots for each species, demonstrating the relationship between geographic and genetic distances ($F_{ST}/1\text{-}F_{ST}$, using the Reich-Patterson $F_{ST}$ estimator) for each pair of sampling sites. Lefthand plots are as in Figure 3. On each righthand plot, the p-value is indicated for a Mantel test of genetic distance as a function of geographic distance; all of these comparisons have a linear model slope not significantly different from 0 and most $F_{ST}$ values between sampling sites are 0.

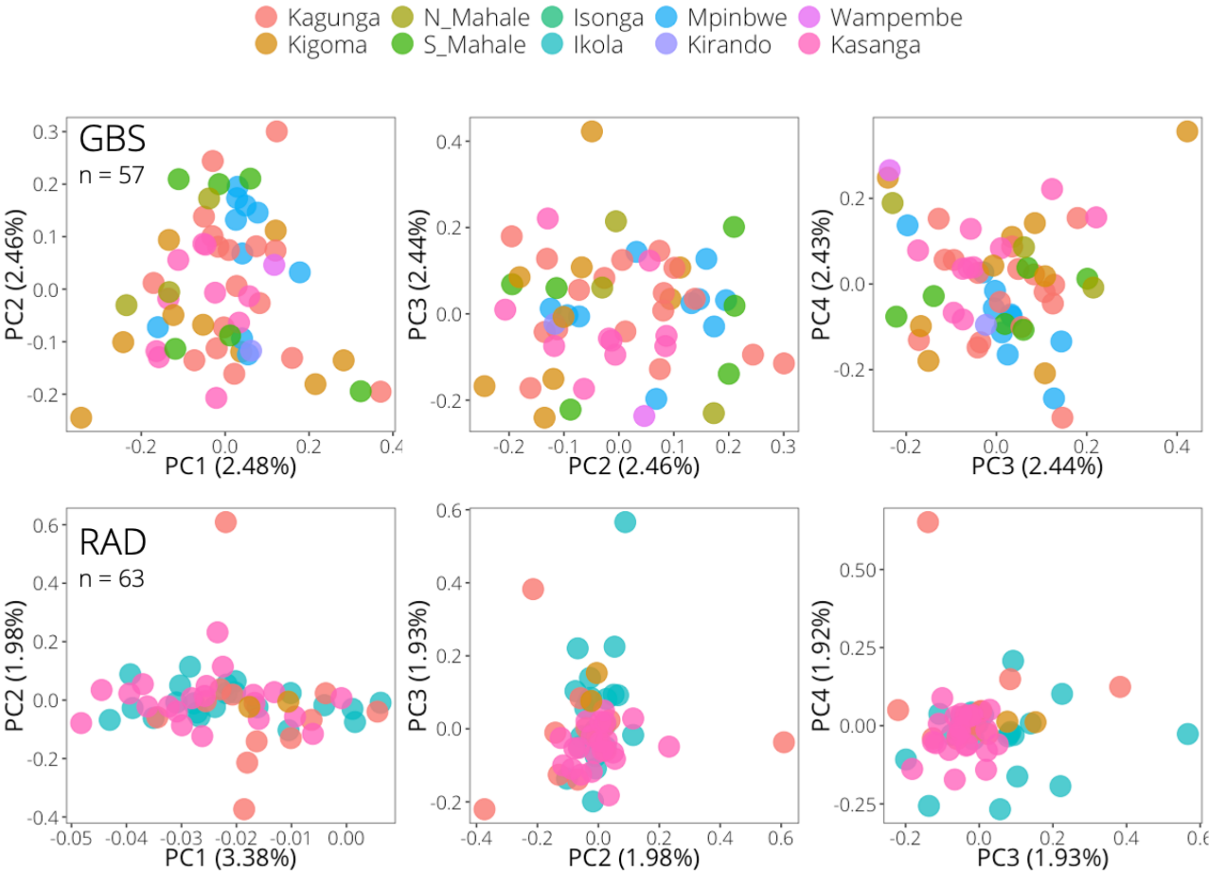


**Figure S4.** Comparison of population structure in *L. stappersii*, as inferred from principal component analysis of genotyping-by-sequencing (GBS) data (top) and restriction-associated digest (RAD) sequencing (bottom). For both data types, the first four PC axes are shown with individuals colored by sampling site. The samples used for GBS were independent from the samples used for the RAD sequencing.

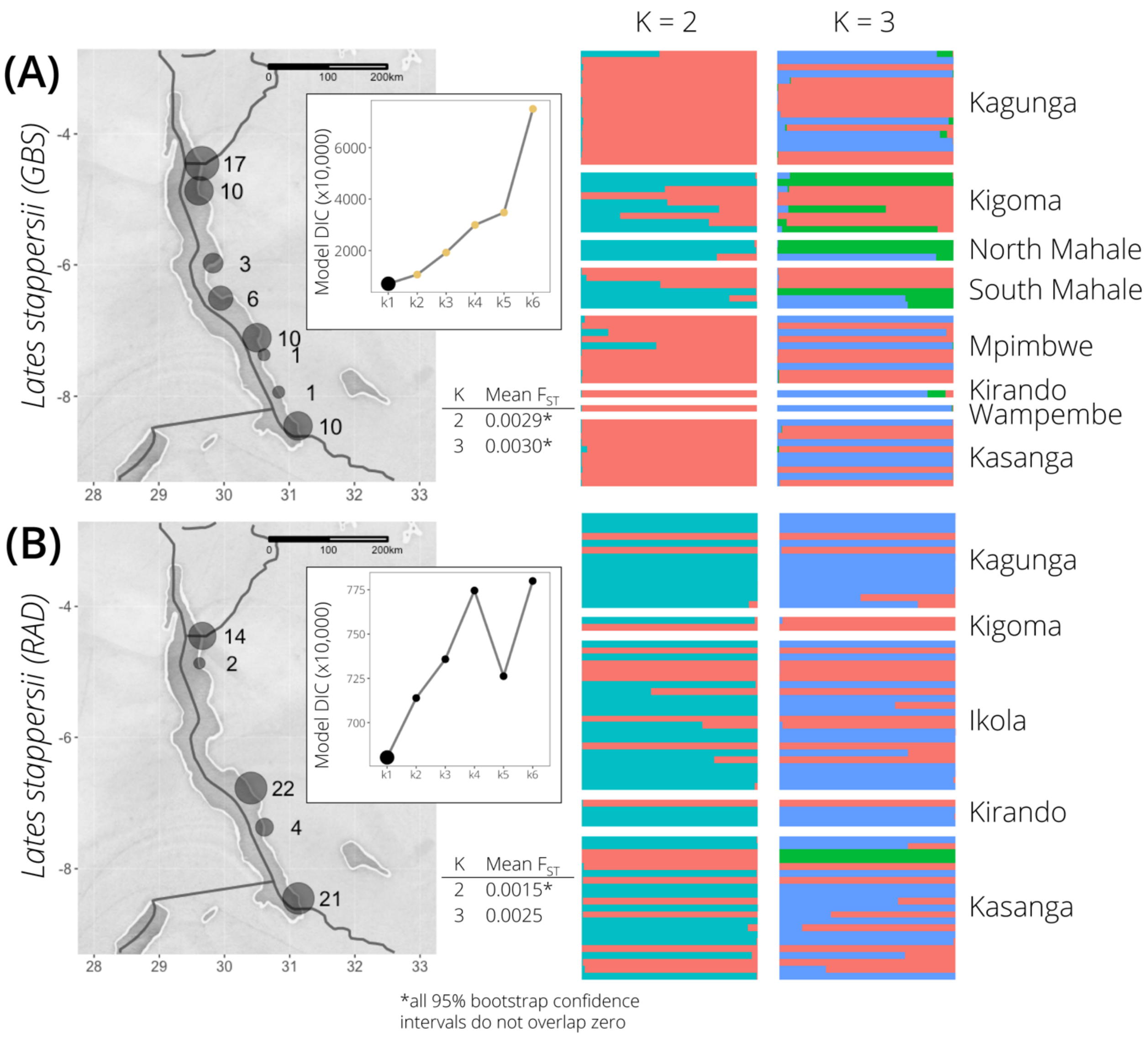


**Figure S5**. Comparison of entropy results at K=2 and K=3 for *L. stappersii* using the (A) GBS and (B) RAD datasets. There are no individuals in common between the GBS and RAD datasets. Sample maps indicate number of individuals collected at each location. Inset plots show DIC values for each value of K from 1 to 6, and mean Reich-Patterson $F_{ST}$ estimates between entropy-identified genetic groups is indicated for K=2 and K=3 ($F_{ST}$ values were not significantly different from 0 at higher levels of K). For calculating $F_{ST}$ estimates, individuals were assigned to groups using a threshold of $q > 0.6$, with individuals with all $q < 0.6$ not having a group assignment.

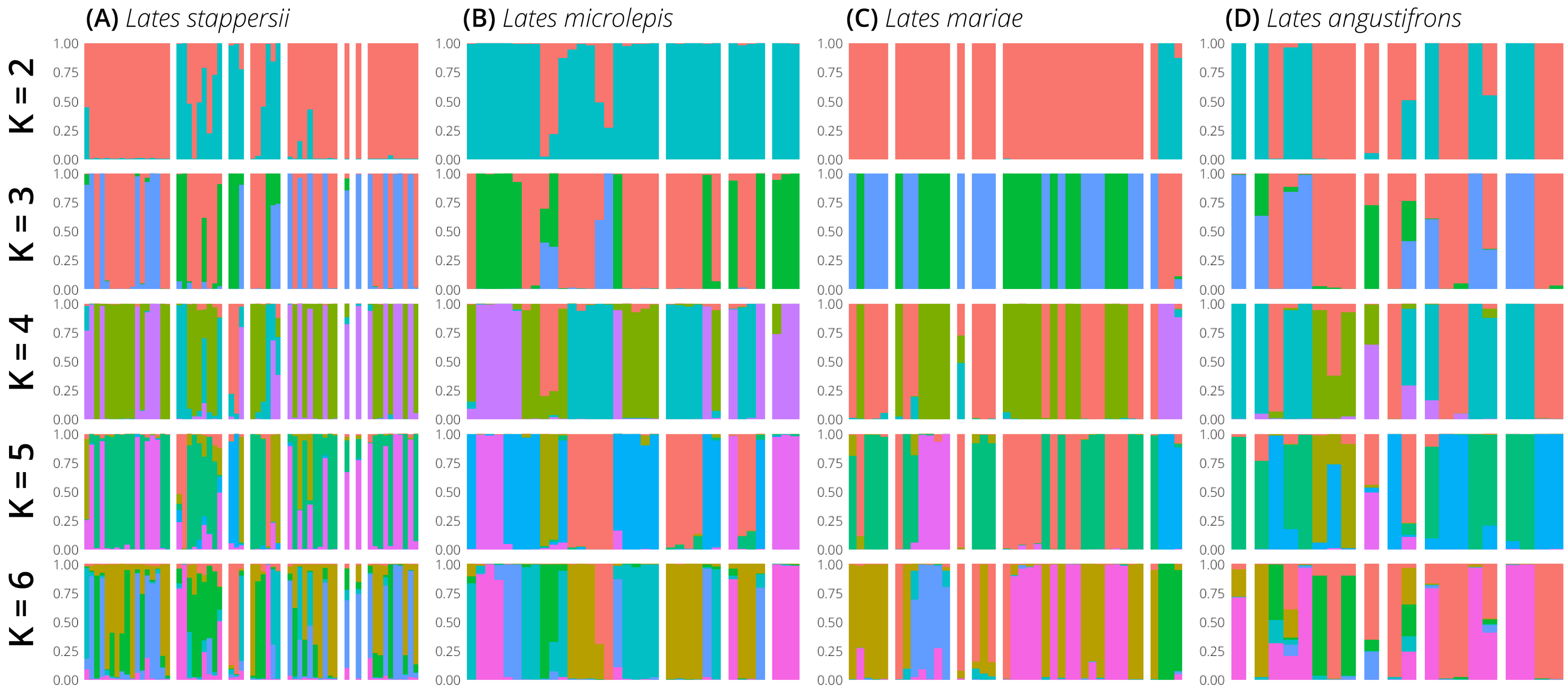


**Figure S6**. Full entropy results from three MCMC chains, from K=2 to K=6, for all four *Lates* species using a missing data cutoff of 50%, minor allele frequency threshold of 0.01, and calling genotypes for sites with a minimum depth of 5. Individuals are sorted according to their collection locations and are in the same location for all values of K. Plots for K=2 and K=3 match those in Fig. 5 in-text.

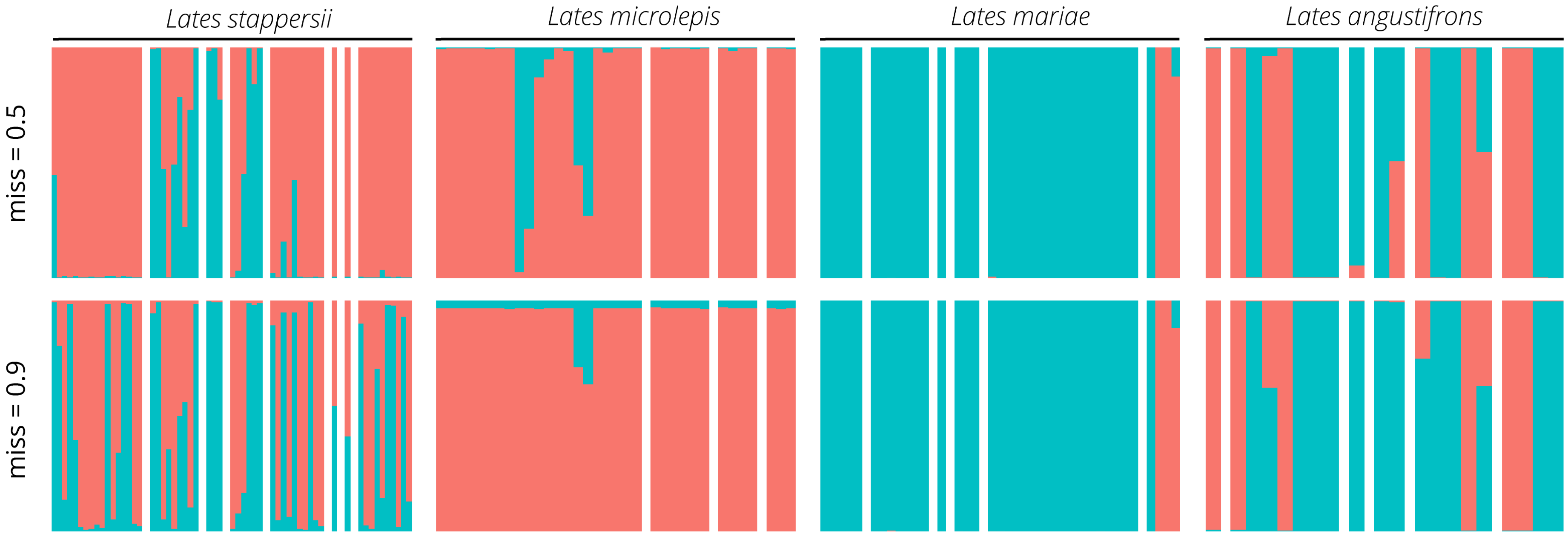


**Figure S7**. Comparison of entropy results at K=2 for datasets with different missing data thresholds, showing results from retaining sites < 50% missing data (top panel, miss = 0.5) and retaining sites with < 10% missing data (bottom panel, miss = 0.9). Individuals are ordered by site, as in Fig. 5 in-text, from north (left) to south (right).

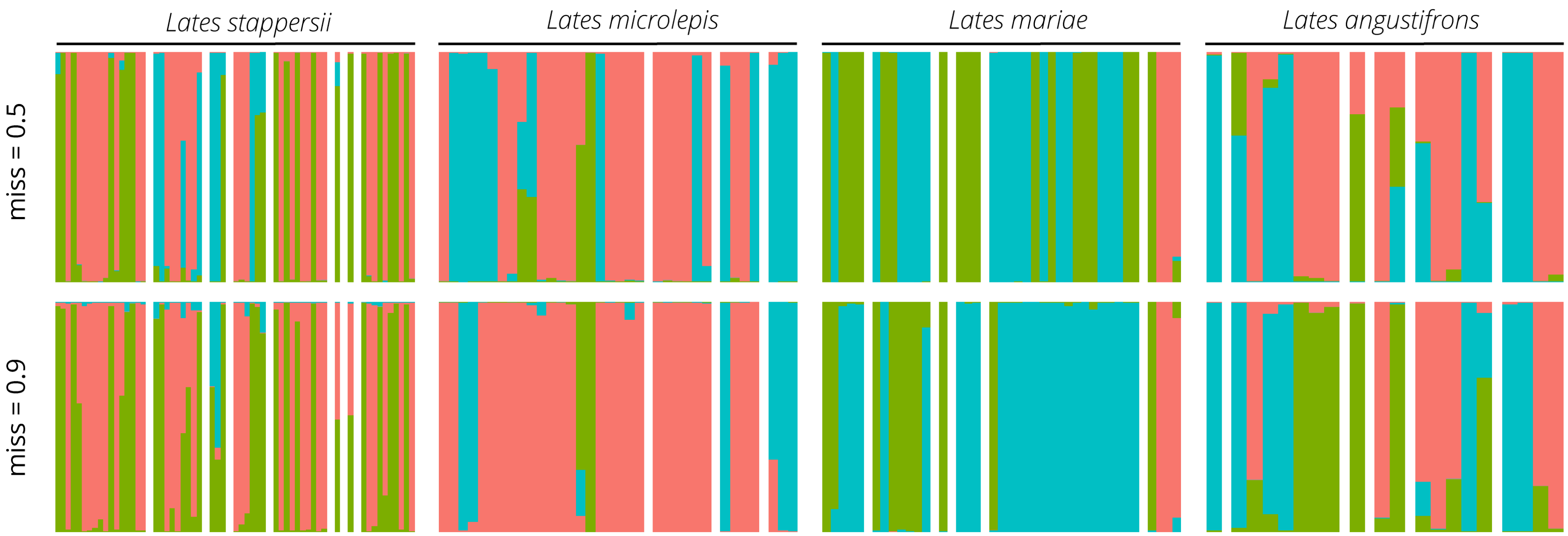


**Figure S8**. Comparison of entropy results at K=3 for datasets with different missing data thresholds, showing results from retaining sites < 50% missing data (top panel, miss = 0.5) and retaining sites with < 10% missing data (bottom panel, miss = 0.9). Individuals are ordered by site, as in Fig. 5 in-text, from north (left) to south (right).

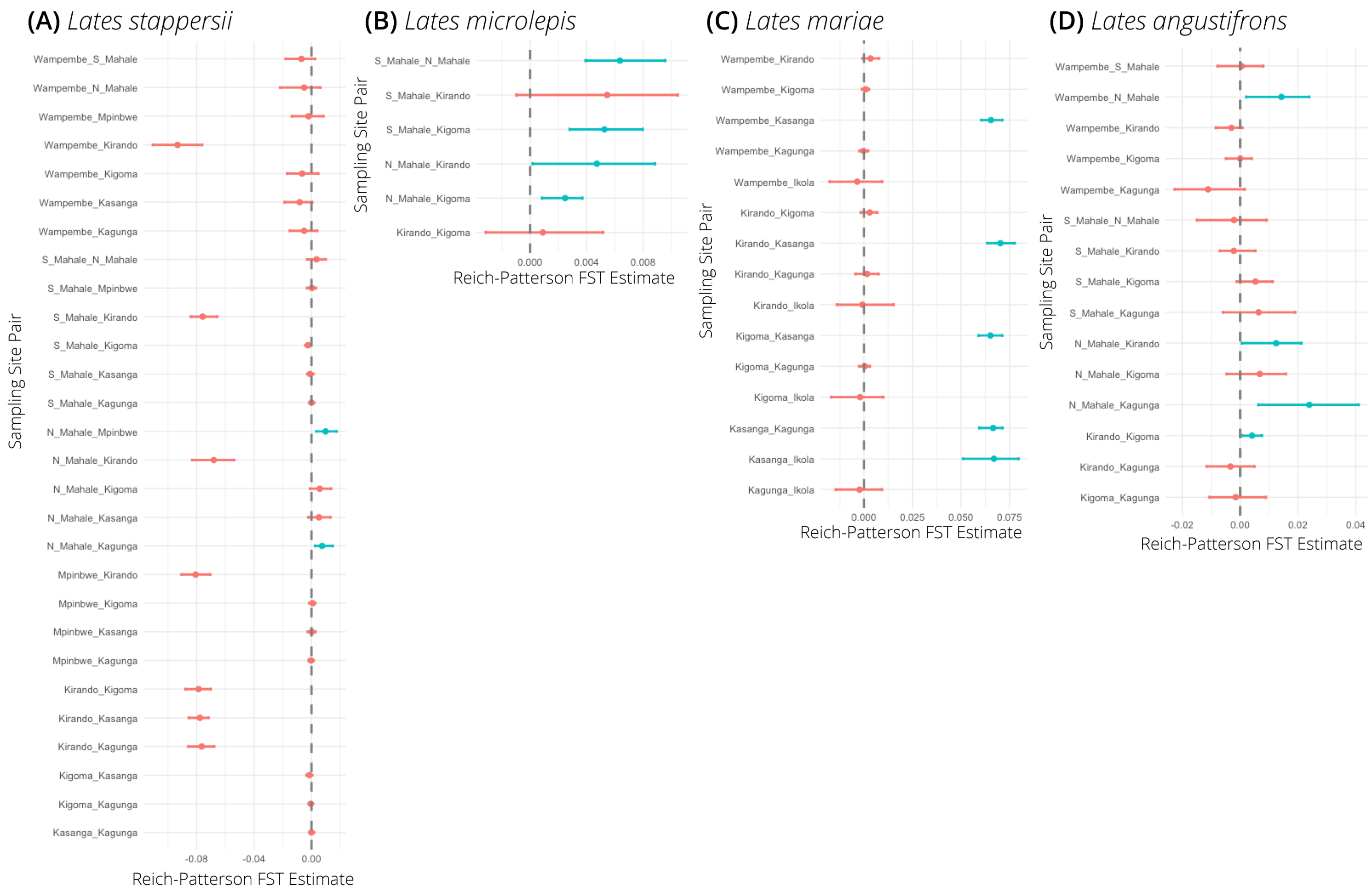


**Figure S9.** Intraspecific Reich-Patterson $F_{ST}$ values between sampling sites. Red indicates values with 95% confidence intervals from 100 bootstraps that overlap 0, while blue indicates values with confidence intervals not overlapping zero. Point estimates of $F_{ST}$s were used in IBD analyses (Figure S3), with negative $F_{ST}$ estimates being corrected to 0.

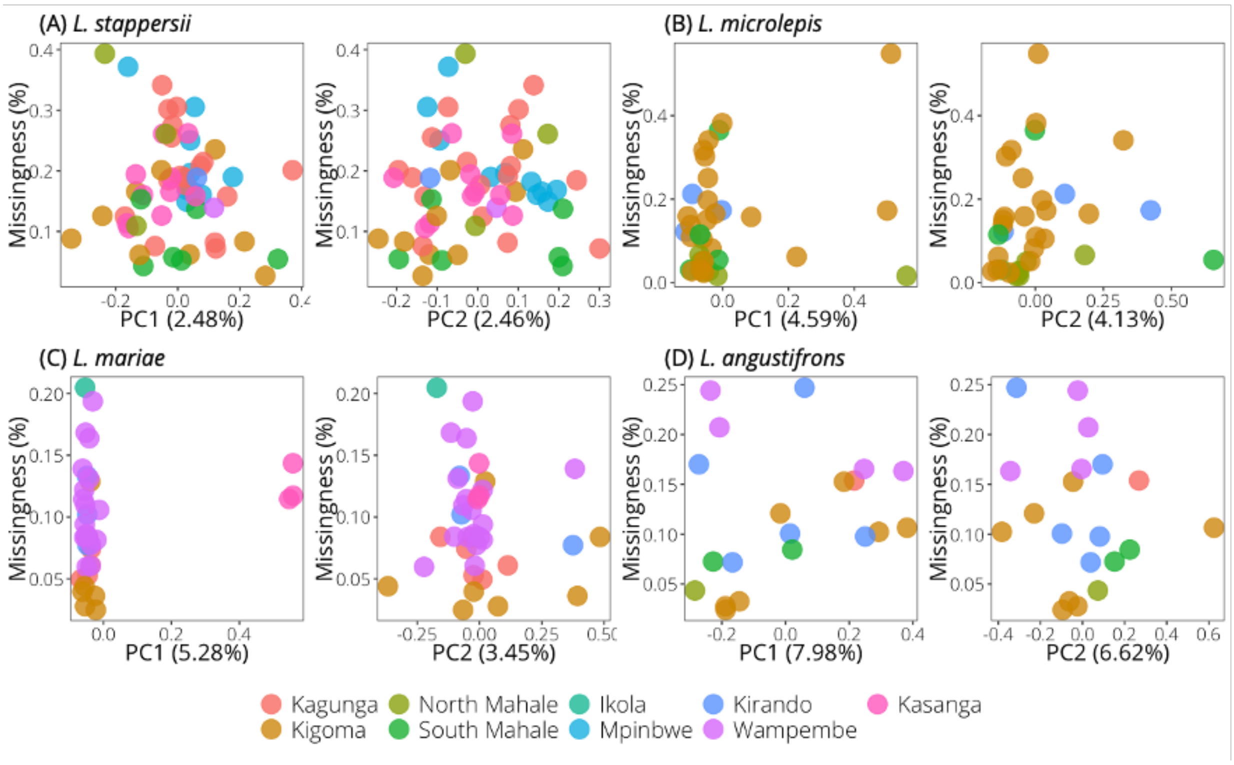


**Figure S10**. Percent missing data in all four species is not correlated with values on PC1 or PC2, suggesting that observed outliers are not simply those with high amounts of missing SNP calls. For each species, the left-hand plot compares PC1 values with percent missing data for each individual, while the righthand plot compares PC2 values with percent missing data. The PC axes match those seen in Figure 3.

91

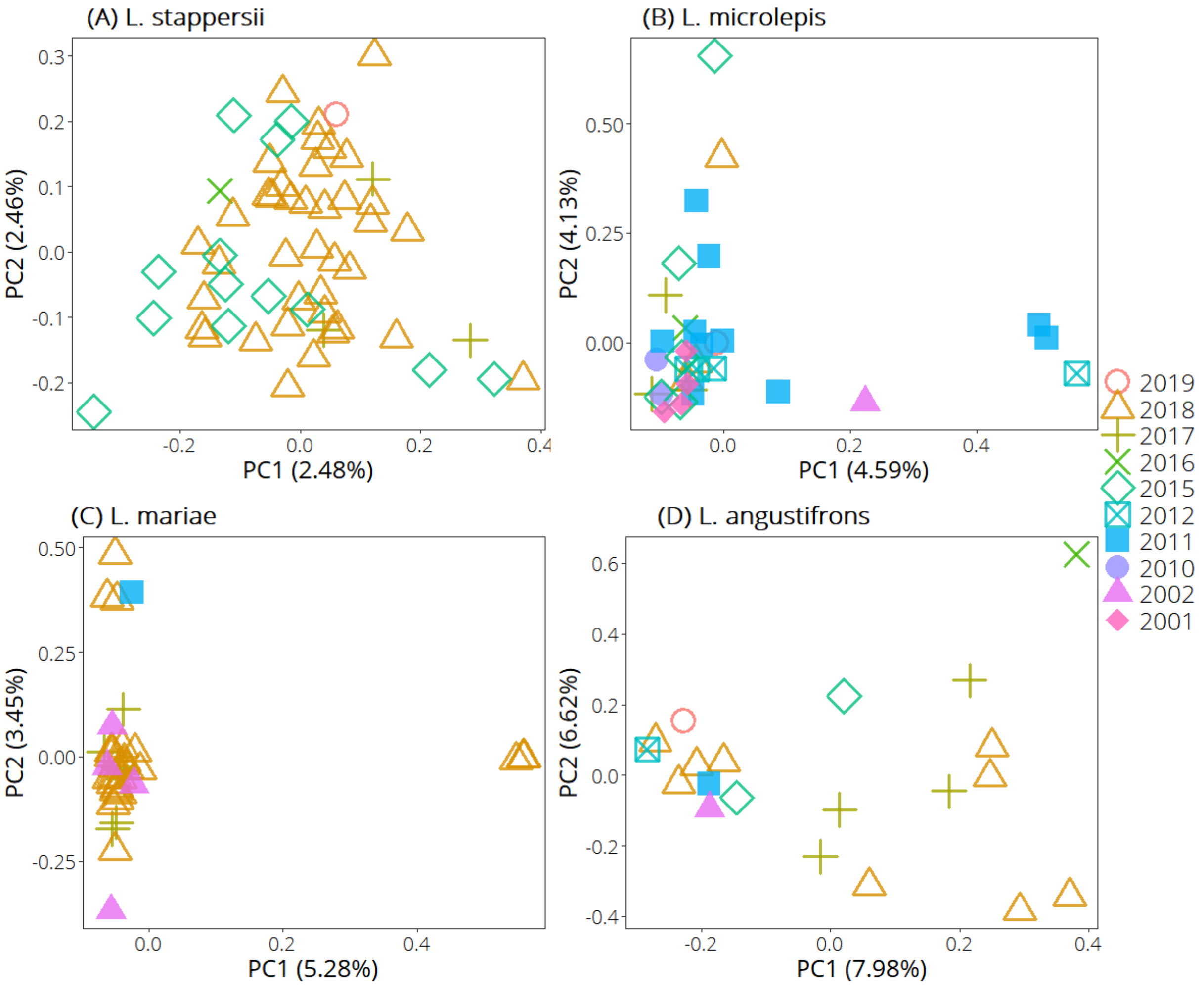


92

93 **Figure S11**. In all four species, collection year is not correlated with values on PC1 or PC2,
94 suggesting that observed outliers are not simply those collected in the same sampling bout.

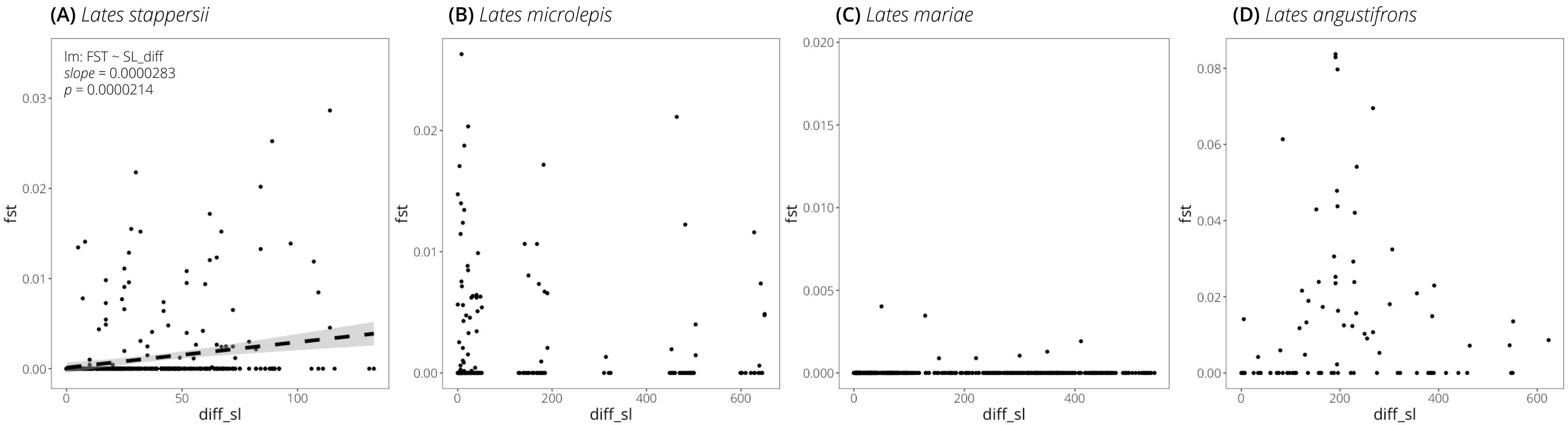


**Figure S12**. Relationship between standard length difference (diff_sl) and pairwise Reich-Patterson FST estimates within species. Linear models were not significant except for in *L. stappersii*, where a weakly positive relationship was observed (i.e., individuals with a greater difference in standard length are more genetically differentiated).

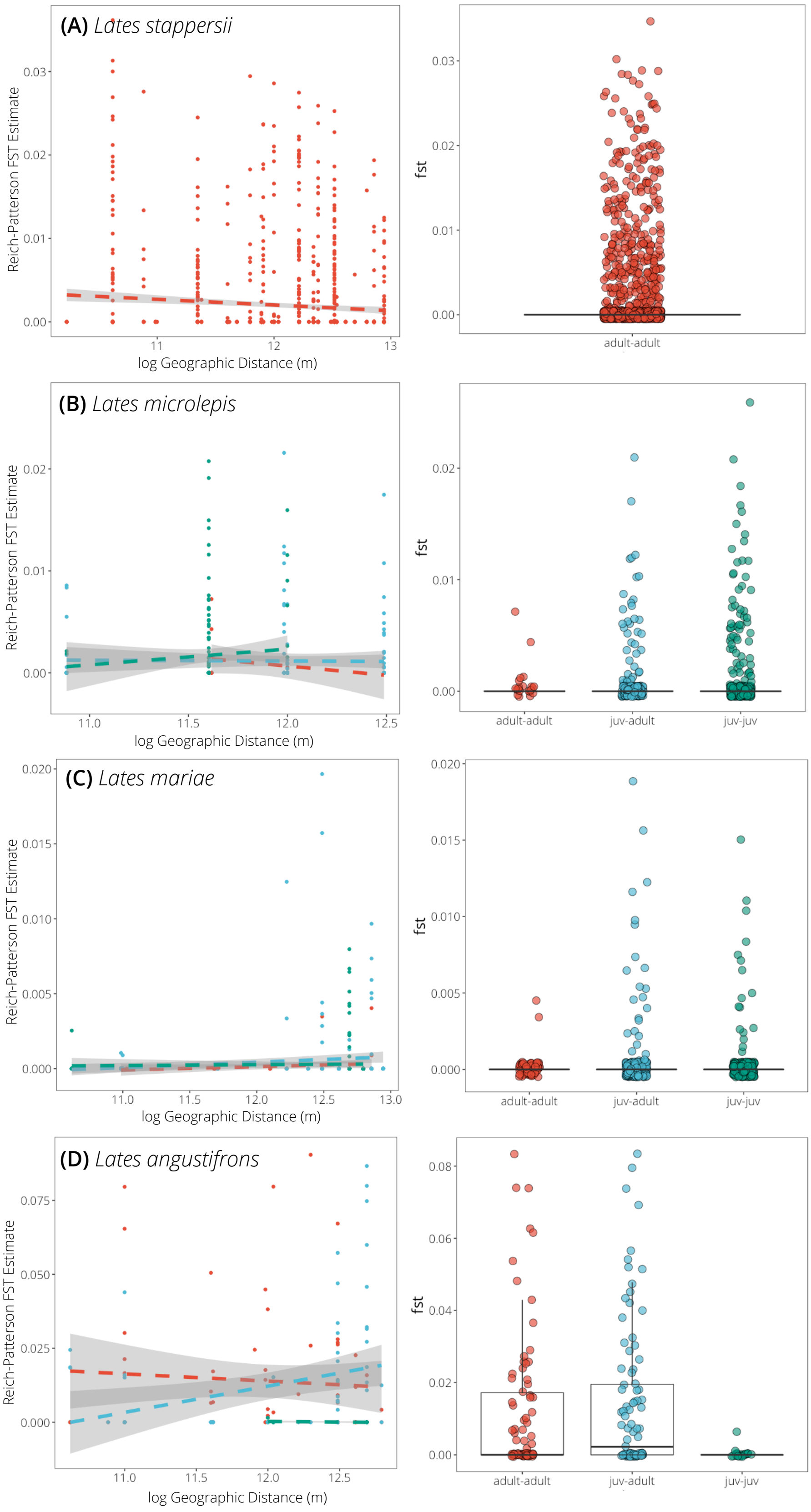


**Figure S13**. Relationship between pairwise genetic and geographic differentiation for each of the four species, grouped by whether both individuals in the comparison are juveniles (juv-juv, green), both are adults (adult-adult, red), or one is a juvenile and one is an adult (juv-adult, blue). All slopes for the linear relationship between genetic and geographic distance (lefthand panels) are not significant, except for juv-adult in *L. angustifrons* and *L. mariae*. Note that there were no *L. stappersii* juveniles caught in our sampling.

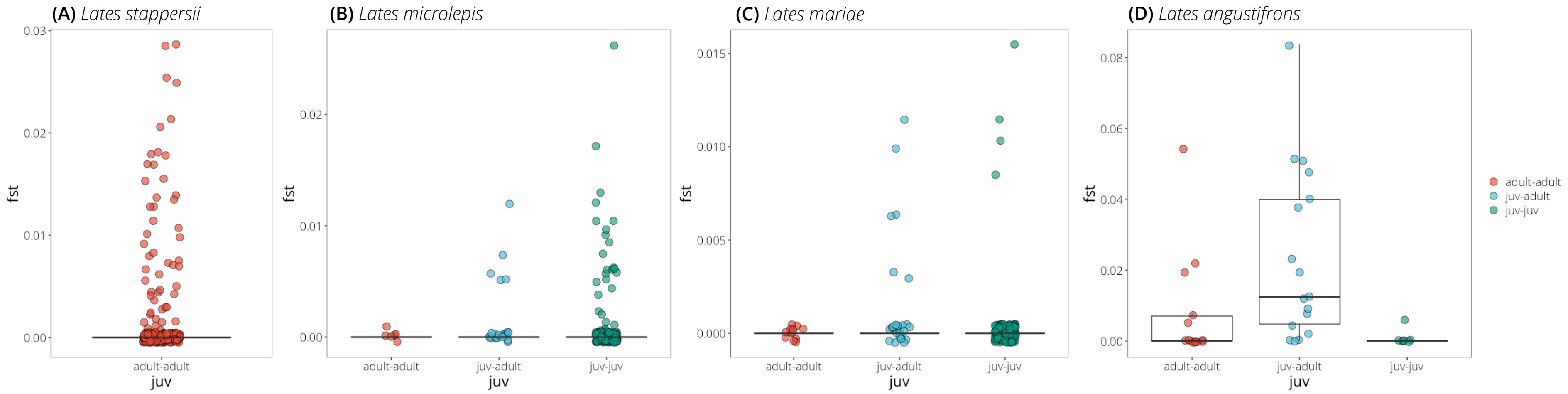


**Figure S14**. Distribution of Reich-Patterson FST values among individuals collected at the *same* sampling site, categorized by whether the comparison is between two adults (adult-adult), two juveniles (juv-juv), or a juvenile and an adult (juv-adult). The only significant difference among categories within species is between the juv-adult category and the adult-adult and juv-juv category in *L. angustifrons*.

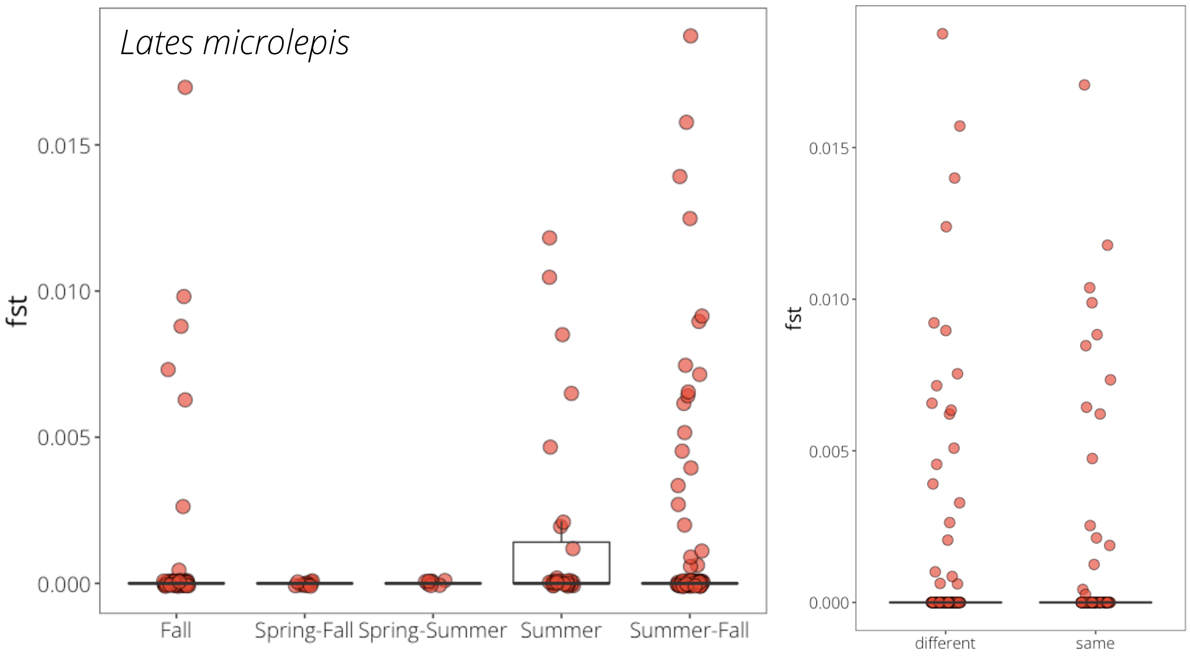


**Figure S15**. Differentiation among individuals caught during the same (Fall, Spring, or Summer) or different times of the year in *L. microlepis*. No differences were found among groups; plots are not shown for *L. stappersii* (due to a lack of juveniles), or *L. angustifrons* and *L. mariae* (due to a lack of differentiation among juveniles).
